# Tubular ERGIC (t-ERGIC): a SURF4-mediated expressway for ER-to-Golgi transport

**DOI:** 10.1101/2021.04.06.438517

**Authors:** Rui Yan, Kun Chen, Ke Xu

## Abstract

The endoplasmic reticulum (ER)-to-Golgi transport is critical to protein secretion and intracellular sorting. Cargo carriers mediating the ER-to-Golgi transport are morphologically diverse, but it remains unclear whether this diversity arises from different cargo receptors, or whether it could lead to differential transport kinetics. Here we report a tubular ER-Golgi intermediate compartment (t-ERGIC) that is induced by the cargo receptor SURF4 and selectively expedites the ER-to-Golgi transport of SURF4 cargoes. Lacking the canonical ERGIC marker ERGIC-53 yet positive for the small GTPase Rab1, the t-ERGIC is further distinct from the stereotypical vesiculo-tubular cluster (VTC) ERGIC by its extremely elongated shape (~10 μm long with <30 nm diameter). With its exceptional surface-to-volume ratio and *en bloc* cargo packaging, high (~2 μm/s) intracellular traveling speeds, and ER-Golgi recycling capability, the t-ERGIC provides an efficient means for trafficking SURF4-bound cargoes. The biogenesis and cargo selectivity of t-ERGIC both depend on SURF4, which recognizes the N-terminus of soluble cargoes and co-clusters with the selected cargoes to expand the ER exit site. At the steady state, the t-ERGIC-mediated fast ER-to-Golgi transport is antagonized by retrograde transport based on the cargo C-terminal ER retrieval signal: we thus demonstrate the fine-tuning of protein trafficking and localization via its primary structure. Together, our results argue that specific cargo-receptor interactions give rise to distinct transport carriers, which in turn regulate the ER-to-Golgi trafficking kinetics.

## INTRODUCTION

About 6,000 human proteins, after synthesized at the endoplasmic reticulum (ER), are transported to the Golgi apparatus for secretion or sorting to other organelles (Barlowe and Helenius, 2016; Dancourt and Barlowe, 2010; Gomez-Navarro and Miller, 2016; Lee et al., 2004). How cells selectively transport different proteins to fulfill their diverse functions has elicited a wealth of research interest. Under current models, ER-to-Golgi trafficking starts at the ER-exit sites (ERESs), where the cargo receptor or the cargo itself recruits the coat protein complex II (COPII) to generate ER-derived vesicles (Brandizzi and Barlowe, 2013; Kurokawa and Nakano, 2019; Lee et al., 2004; Zanetti et al., 2012). Whereas in yeasts these vesicles appear to directly reach the Golgi apparatus via cytoplasmic diffusion (Kurokawa and Nakano, 2019; Lee et al., 2004), in animal cells they relay the cargo to the ER-Golgi intermediate compartment (ERGIC) for microtubule-dependent sorting to the Golgi apparatus (Appenzeller-Herzog and Hauri, 2006; Saraste and Marie, 2018). Additionally, proteins lacking affinity to any receptors may be transported via “bulk flow” through nonspecific packaging into the ERES (Barlowe and Helenius, 2016; Lee et al., 2004).

Contrasting with the diversity of protein cargoes and receptors identified over the past decades, the ERGIC is often simply represented by the presence of the membrane lectin ERGIC-53/LMAN1, which may not define all carriers mediating ER-to-Golgi trafficking (Appenzeller-Herzog and Hauri, 2006; Hauri and Schweizer, 1992; Saraste and Marie, 2018). Whereas electron microscopy often shows ERGIC as vesiculo-tubular clusters (VTCs) <~1 μm in size (Saraste and Svensson, 1991; Schweizer et al., 1988), elongated tubular carriers have also been noted (Bannykh et al., 1996; Klumperman et al., 1998; Mironov et al., 2003). Live-cell fluorescence microscopy using synchronized cargoes has also occasionally noted long (>2 μm) tubular carriers in ER-to-Golgi trafficking (Ben-Tekaya et al., 2005; Blum et al., 2000; Marra et al., 2001; Presley et al., 1997; Shomron et al., 2019; Simpson et al., 2005), yet some cargoes do not seem to access this pathway (Boncompain et al., 2012; Westrate et al., 2020). Whether these morphologically distinct carriers are specific to different protein cargoes, and if so, what determines such selectivity, remain elusive. Furthermore, the functional significance of this morphological diversity in regulating transport kinetics remains unexplored.

Here we report tubular ERGIC (t-ERGIC), a distinct class of ERGIC that specifically expedites the ER-to-Golgi transport of soluble cargoes of the receptor SURF4, the mammalian homolog of the yeast cargo receptor Erv29p (Belden and Barlowe, 2001; Dancourt and Barlowe, 2010; Mitrovic et al., 2008). The t-ERGIC lacks ERGIC-53 but is marked by the small GTPase Rab1, and further differs from the canonical ERGIC/VTCs by its extremely slender shape. After *de novo* generation at expanded ERESs via SURF4-cargo interactions, the t-ERGIC travels and recycles in the cell at high speeds to enable efficient ER-to-Golgi transport.

## RESULTS

### Identification of a Highly Elongated Tubular Organelle through Mislocalized DsRed2-ER-5

We initially attempted to deliver the red fluorescent protein (FP) DsRed2 into the ER lumen of COS-7 cells via an N-terminal signal peptide. Curiously, whereas DsRed2-ER-3 (Figure 1A) correctly localized to the ER (Figure 1B), a similar construct with slightly varied linkers, DsRed2-ER-5 (Figure 1A), only showed up the expected ER localization in ~20% of the transfected cells. A preponderance (~45%) of the cells had fluorescence mainly distributed in peculiar, 2-20 μm long tubular organelles (TOs) (Figures 1B-1D), while another major population (~30%) had most fluorescence in small vesicles (Figures 1D, S1A).

**Figure 1.**
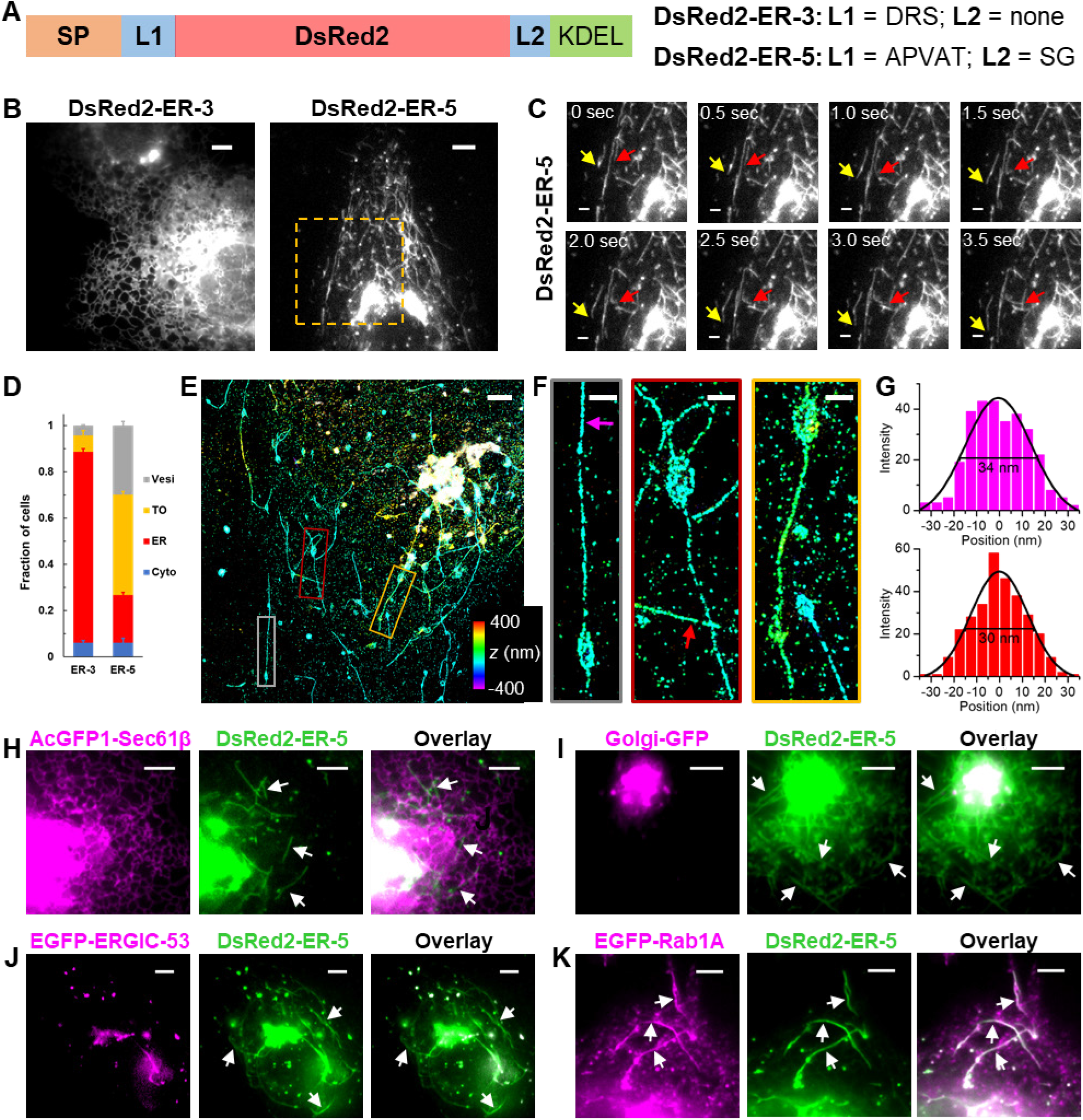
DsRed2-ER-5 Mislocalizes to an Extremely Slender, Rab1-Positive Tubular Organelle. (A) Schematics of the sequences of DsRed2-ER-3 and DsRed2-ER-5. SP: signal peptide; L: linker. (B) Representative live-cell images of DsRed2-ER-3 and DsRed2-ER-5 expressed in COS-7 cells. (C) Time-lapse series of the boxed region in (B), showing the morphology and fast motion of the tubular organelles (TOs) (arrows). See also Video S1. (D) Classification of the dominating distribution modes of DsRed2-ER-3 and DsRed2-ER-5 in each cell: ER, TO, vesicles (Vesi), and cytoplasm (Cyto). Error bars: SEM (n = 4 with ~50 cells in each replicate). (E) 3D-STORM image of immunolabeled DsRed2-ER-5 in a COS-7 cell. Colors encode axial positions. (F) Close-ups of the TOs in the three colored boxes in (E). (G) STORM intensity profiles across the widths of two TOs at the magenta and red arrows in (F). Black curves: Gaussian fits with FWHM (full width at half maximum) of 34 and 30 nm, respectively. (H-K) Dual-color live-cell images of DsRed2-ER-5 (green) with the ER marker AcGFP1-Sec61β (H), the Golgi marker Golgi-GFP (I), the canonical ERGIC marker EGFP-ERGIC-53 (J), and the small GTPase EGFP-Rab1A (K). Arrows point to TOs. Scale bars: 5 μm (B,H-K); 2 μm (C,E); 500 nm (F). See also Figure S1.

The DsRed2-ER-5-containing TOs each had one or a few enlarged vesicular bodies and a thin projection, and underwent frequent deformation, fission, and fusion as they moved actively in the cell at typical speeds of 1-2 μm/s (Figure 1C, Video S1). The vesicular bodies were often found at the tubule ends, yet they also frequently slid along the TOs, indicating high structural flexibility (Video S1). Three-dimensional (3D) STORM super-resolution microscopy (Huang et al., 2008) further revealed their unusual ultrastructure (Figures 1E and 1F): Whereas the vesicular bodies were spheroids ~500 nm in size, the elongated projections were extremely thin with apparent diameters of ~30 nm (Figure 1G). Given the ~20 nm resolution of STORM and the immunolabeling antibody sizes (Huang et al., 2008), the true diameters should thus be <30 nm. Similar TOs were also found in U2OS and HeLa cells (Figures S1B and S1C).

To identify these TOs, we performed dual-color fluorescence microscopy of DsRed2-ER-5 and different organelle markers in living cells. The TOs did not have soluble or membrane markers of the ER (Figures 1H, S1D), nor were they positive for common markers of endosomes, autophagosomes, or lysosomes (Figures S1E-S1I). Instead, we noticed dynamic interactions of the TOs with the Golgi apparatus, where the FP accumulated (Figure 1I, Video S1), suggesting its involvement in the secretory pathway. Interestingly, whereas the canonical ERGIC marker ERGIC-53 colocalized with DsRed2-ER-5 in vesicle-like structures, it was absent from the TOs (Figure 1J). However, when ERGIC-53 was overexpressed at high levels, it also entered a subset of the TOs (Figure S1J), implying interactions and material exchange between the TOs and the canonical ERGIC. COPII and COPI coats also only partly colocalized with DsRed2-ER-5 in puncta but not in the TOs (Figures S1K and S1L).

We next found Rab1A/B, the major small GTPases in ER-to-Golgi trafficking (Plutner et al., 1991; Stenmark, 2009; Tisdale et al., 1992), are enriched on the surface of the DsRed2-ER-5-containing TOs, a result supported by both expressed EGFP-Rab1A (Figures 1K, S1M) and the immunostaining of endogenous Rab1A and Rab1B (Figures S1N and S1O). Time-lapse imaging captured the dynamic flow of DsRed2-ER-5 from the ER to small vesicles (Figure S1P). In this process, the TOs only existed before DsRed2-ER-5 reached its maximal signal at the Golgi, suggesting that they are intermediates between the ER and the Golgi. Together, we identified the DsRed2-ER-5-containing, highly elongated TO as a Rab1-coated ERGIC that is not normally enriched with ERGIC-53. Hereafter we refer to it as “tubular ERGIC (t-ERGIC)”.

Another construct, GCaMP6s-ER-5, exhibited a similar phenotype as DsRed2-ER-5, which allowed us to image the t-ERGIC with a live-cell microtubule marker 2xmCherry-EMTB, thus showing the t-ERGICs traveling along microtubules (Video S2).

### The t-ERGIC Mediates ER-to-Golgi Trafficking, and is Formed through Both *De Novo* Generation and Fusion

To understand how DsRed2-ER-5 escaped from the ER, we examined the effects of different small molecules that respectively inhibited ER-to-Golgi transport (brefeldin A), induced ER stress (thapsigargin and dithiothreitol), and inhibited ER-associated degradation (CB-5083 and MG132). Flow cytometry showed that only brefeldin A significantly increased the intracellular DsRed2 fluorescence (Figure S2A). The time-dependent increase of intracellular retention with brefeldin A treatment (Figures 2A and 2B) was accompanied by a redistribution of the DsRed2-ER-5 fluorescence back to the ER (Figures 2C, S2B). These results suggest that DsRed2-ER-5 is targeted to the ER-to-Golgi transport pathway through t-ERGIC.

**Figure 2.**
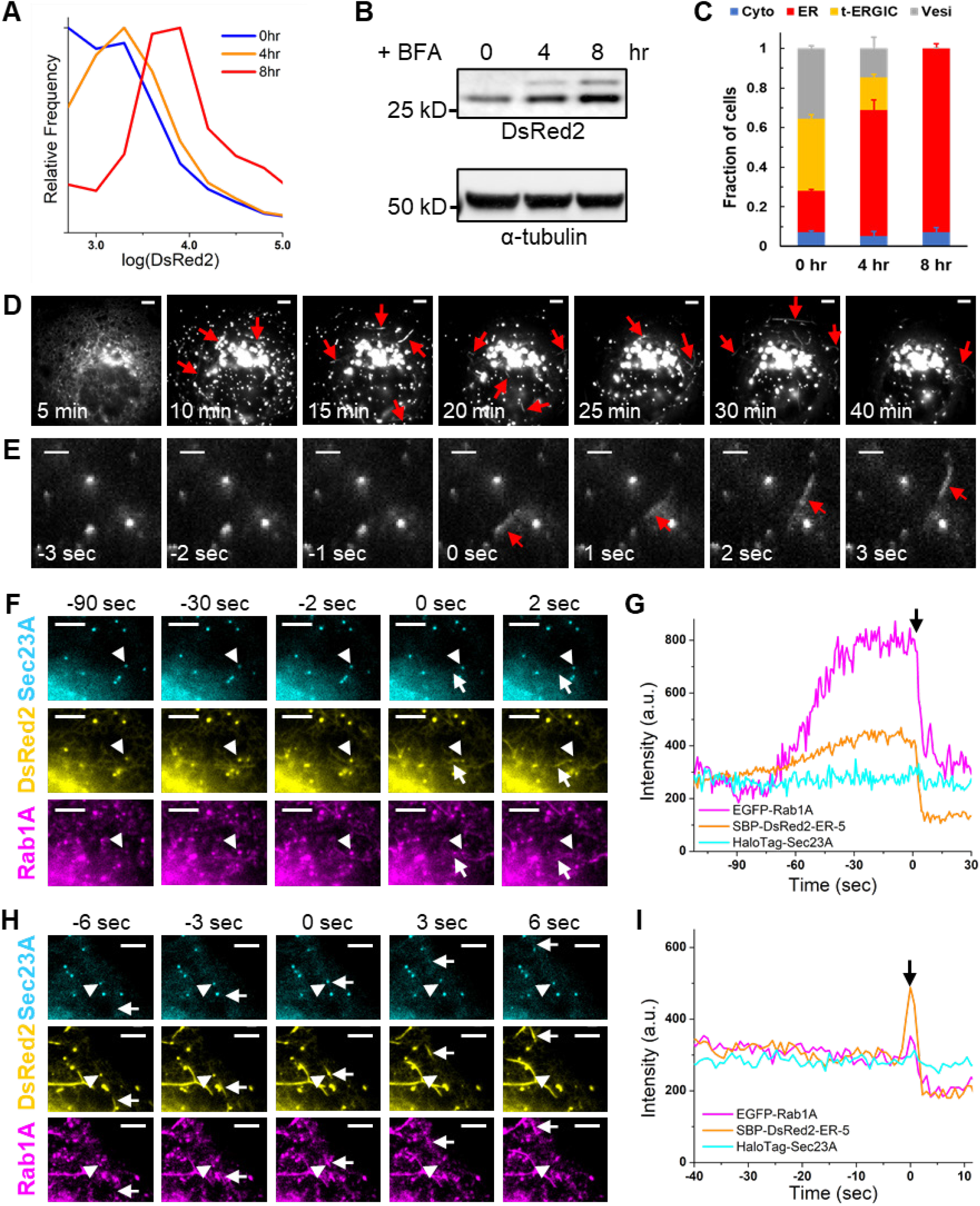
The t-ERGIC Mediates ER-to-Golgi Trafficking, and is Formed through Both *De Novo* Generation and Fusion. (A-C) Flow cytometry histograms (A), lysate immunoblots (B), and subcellular distribution (C) of DsRed2-ER-5 for transfected COS-7 cells treated with 1 μM brefeldin A (BFA) for 0, 4, and 8 hr. Error bars: SEM (n = 3 with ~50 cells in each replicate). (D) Representative RUSH image sequence of SBP-DsRed2-ER-5 after the addition of 80 μM biotin at time 0. Arrows indicate t-ERGICs. See also Video S3. (E) *De novo* generation of SBP-DsRed2-ER-5-positive t-ERGIC (arrow) in RUSH. Time 0 corresponds to when budding occurred. Biotin was added at −5 min for cargo release. (F) Image sequences of EGFP-Rab1A, SBP-DsRed2-ER-5, and JF635-labeled HaloTag-Sec23A in a RUSH experiment showing the *de novo* generation of a t-ERGIC (arrow) from the ERES (arrowhead). Time 0 corresponds to when budding occurred. Biotin addition corresponded to −20 min. (G) Fluorescence intensity time traces of the three color channels for the ERES indicated by the arrowhead in (F). (H) Another image sequence of an ERES (arrowhead) in the same RUSH experiment as in (F), showing its fusion with a pre-existing t-ERGIC (arrow). Time 0 corresponds to when fusion occurred. Biotin addition corresponded to −28 min. (I) Fluorescence intensity time traces of the three color channels for the ERES indicated by the arrowhead in (H). Scale bars: 5 μm (D,F,H); 2 μm (E). See also Figure S2.

To elucidate the biogenesis and potential functions of the t-ERGIC, we next employed the Retention Using Selective Hooks (RUSH) assay (Boncompain et al., 2012) to synchronize the release of DsRed2-ER-5 from the ER. In RUSH, the cargo is tagged with a streptavidin binding peptide (SBP) and thus initially retained in the ER by an ER-resident streptavidin “hook”. The addition of biotin outcompetes SBP for streptavidin binding and so enables the synchronized onset of ER-to-Golgi transport of the cargo. Insertion of the SBP tag at the N-terminus of DsRed2-ER-5 (Figure S2C) did not alter the phenotype (Figures S2D and S2E). The SBP-DsRed2-ER-5 cargo was then co-expressed with a streptavidin-KDEL hook, which retained most of the fluorescence signal in the ER two days after transfection. Upon release of the cargo by biotin, we observed that the cargo first concentrated at the ERES and vesicle-like structures (Figure 2D, Video S3). The t-ERGIC emerged right after ER exit (Figures 2D and 2E; Video S3). Single-particle tracking of the vesicles and TOs showed an overall centripetal movement, as the fluorescence redistributed from the peripheral ER to the Golgi apparatus (Figures S2F and S2G). These results confirm that the t-ERGIC mediates anterograde ER-to-Golgi transport.

A closer examination of the RUSH image sequences unveiled two modes of biogenesis for t-ERGIC: *de novo* formation vs. elongation of existing t-ERGIC through fusion. For the first mode, three-color live imaging of the cargo with the ERES marker Sec23A and the t-ERGIC marker Rab1A (Figures 2F and 2G, S2H) showed that the cargo was first enriched at the ERES, where Rab1A gradually accumulated, leveled off, and then budded off together with the cargo into newly formed t-ERGIC tubules. The budded t-ERGIC then separated from the ERES, leaving behind the Sec23A COPII coat (Figures 2F, S2H), from which another t-ERGIC could bud again (Figure S2I). Accordingly, local fluorescence intensity time traces showed that DsRed2-ER-5 and Rab1A both accumulated at the ERES before they simultaneously budded into the t-ERGIC in a single step, whereas Sec23A stayed constant (Figure 2G). In the second mode, existing t-ERGIC tubules actively collected more cargo at ERES as they rapidly traversed the cell (Figures 2H and 2I, Video S3). In particular, we often noticed cases in which tubules generated from the Golgi traveled retrogradely to fuse with the ERES and bring more cargo back to the Golgi (Figure S2J, Video S3), a cycling behavior that has been noted previously for ERGIC (Ben-Tekaya et al., 2005; Marra et al., 2001; Sannerud et al., 2006). Thus, the t-ERGIC is a carrier organelle that buds from the ERES and shuttles between the ER and the Golgi to mediate the anterograde transport of cargo proteins.

The accumulation of Rab1A at the ERES before t-ERGIC formation prompted us to test its role in t-ERGIC biogenesis. Using a dominant negative Rab1A mutant (N124I) (Moyer et al., 2001; Westrate et al., 2020), we observed that the transfected cells were unable to generate t-ERGIC tubules after the cargo accumulated at the ERES (Figure S2K). Together, our RUSH assays indicate that the Rab1A-dependent t-ERGIC mediates the ER-to-Golgi trafficking of DsRed2-ER-5.

### Fast ER-to-Golgi Trafficking via the t-ERGIC Is Determined by the N-termini of Soluble Cargoes

Our unexpected discovery of t-ERGIC through DsRed2-ER-5 raises the question of why a similar construct, DsRed2-ER-3, localized predominantly in the ER (Figures 1B, 1D). Given the small dissimilarities between the two constructs (Figure 1A), we wondered whether the property of the N-terminus after the signal-peptide cleavage (P1’) could be important to the fate of the cargo protein. The N-terminus of the cleaved DsRed2-ER-5 begins with the hydrophobic APV tripeptide, whereas that of DsRed2-ER-3 starts with the charged DRS (Figure 1A). Consequently, we constructed a point mutant of DsRed2-ER-5, where the P1’ alanine (A) was substituted by glutamic acid (E). Remarkably, this EPV-DsRed2-ER-5 variant (Figure 3A) phenocopied DsRed2-ER-3 (Figures 3B and 3C), thus a good control for the original DsRed2-ER-5 (hereafter APV-DsRed2-ER-5) for our mechanistic investigations.

**Figure 3.**
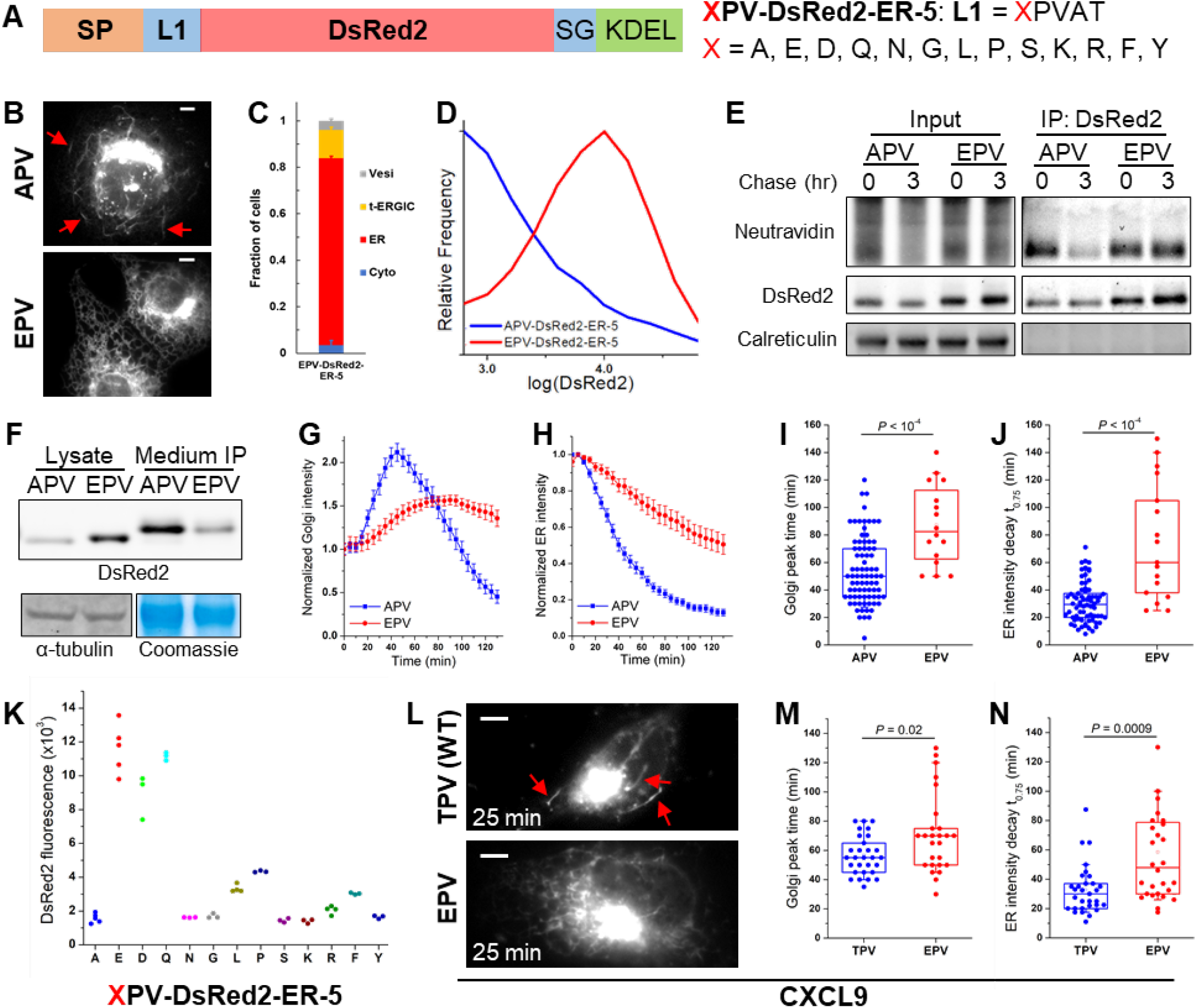
The N-terminus of the Cargo Determines Its Transport with the t-ERGIC and ER-to-Golgi Trafficking Efficiency. (A) Sequences of the XPV-DsRed2-ER-5 mutations we examined, with varied N-termini after the signal peptide (SP). The original DsRed2-ER-5 has X=A (APV-DsRed2-ER-5). (B) Representative fluorescence micrographs of APV/EPV-DsRed2-ER-5 in COS-7 cells. Arrows point to t-ERGICs. (C) Subcellular distribution of EPV-DsRed2-ER-5. Error bars: SEM (n = 3 with ~50 cells in each replicate). (D) Flow cytometry histograms of APV/EPV-DsRed2-ER-5. (E) Azidohomoalanine-biotin-alkyne pulse-chase of APV/EPV-DsRed2-ER-5. Newly synthesized proteins were labeled by azidohomoalanine click chemistry and detected by NeutrAvidin (see Methods). (F) Immunoblots of intracellular (cell lysate) and secreted (anti-FLAG immunoprecipitation from the culture medium) APV/EPV-FLAG-DsRed2-ER-5. (G,H) Golgi (G) and peripheral ER (H) fluorescence intensity time traces of APV/EPV-SBP-DsRed2-ER-5 in RUSH, pooled from 70 cells from 5 independent runs (APV) or 17 cells from 3 independent runs (EPV). Error bars: SEM. 80 μM biotin was added at time 0. (I,J) Comparison of the time to the peak fluorescence in the Golgi (I) and the time of fluorescence decay to 75% of the start in the ER (J) of APV/EPV-SBP-DsRed2-ER-5 in RUSH. Whiskers and boxes show 10%, 25%, 50%, 75%, and 90% quantiles. (K) Median intracellular fluorescence of different XPV-DsRed2-ER-5 variants expressed in COS-7 cells, as determined by flow cytometry. (L) Representative fluorescence micrographs of TPV/EPV-CXCL9-mCherry-SBP in RUSH. 80 μM biotin was added at time 0. Arrows point to t-ERGICs. See also Video S4. (M,N) Comparison of the time to the peak fluorescence in the Golgi (M) and the time of fluorescence decay to 75% of the start in the ER (N) of TPV/EPV-CXCL9-mCherry-SBP in RUSH. Whiskers and boxes show 10%, 25%, 50%, 75%, and 90% quantiles. Scale bars: 5 μm. *P* values are calculated by two-tailed *t* test. See also Figure S3.

Flow cytometry of COS-7 cells transfected with APV-DsRed2-ER-5 and EPV-DsRed2-ER-5 showed markedly higher intracellular fluorescence for the latter (Figure 3D). Pulse-chase experiments indicated that the APV version was selectively removed from the cell (Figure 3E). With FLAG-tagged versions of APV/EPV-DsRed2-ER-5 (Figure S3A), which, without altering the APV/EPV phenotypes (Figures S3B and S3C), facilitated immunoprecipitation from the culture medium, we next found substantially higher extracellular secretion and lower intracellular retention for the APV version (Figure 3F). Immunoblots of ER stress indicators indicated no noticeable activation (Figure S3D), suggesting that the different fates of the APV and EPV variants were attributed to physiological secretory pathways. Similar contrasting behavior of the two variants was observed in U2OS and HeLa cells (Figure S3E), as well as for APV/EPV variants of the GCaMP6s FP (Figures S3F and S3G).

RUSH experiments showed that contrasting the fast, t-ERGIC-mediated ER-to-Golgi transport of APV-SBP-DsRed2-ER-5 (Figure 2D), EPV-SBP-DsRed2-ER-5 (Figure S3A) did not enter t-ERGIC and was slowly transported to the Golgi after biotin release (Figure S3H). Quantification of the fluorescence intensity in the Golgi area showed a substantially faster rise for the APV version after cargo release (Figures 3G and 3I). Accordingly, fluorescence in the ER decayed significantly faster in the APV-transfected cells (Figures 3H and 3J). These results demonstrate dramatic differences in the ER-to-Golgi transport pathway and efficiency between APV-and EPV-DsRed2-ER-5, which explain their different intracellular retentions at the steady state.

To further examine the effects of the cargo N-terminus, we compared 13 different amino acid residues at the P1’ position (Figure 3A). Interestingly, we found DsRed2 to be strongly retained in the ER when glutamic acid (E), aspartic acid (D), or glutamine (Q) was present at the P1’ position, but often entered the t-ERGIC and got exported out of the ER when the P1’ position was other amino acids, including the structurally similar asparagine (N) (Figures 3K, S3I and S3J).

To test whether this “N-terminus rule” for t-ERGIC-mediated ER export is relevant to endogenous proteins, we examined a normally secreted cytokine CXCL9, which has a native TPV P1’ N-terminus. RUSH showed that the wild-type CXCL9 was transported *via* the t-ERGIC, whereas few t-ERGICs were involved in the trafficking of a mutant with an EPV N-terminus (Figure 3L, Video S4). Concomitantly, substantially faster ER-to-Golgi transport was found for the former (Figures 3M and 3N).

Collectively, our data indicate that a group of cargoes with explicit N-terminal features are routed to the t-ERGIC-mediated fast ER-to-Golgi trafficking pathway.

### The Biogenesis and Cargo Selectivity of t-ERGIC Both Depend on SURF4

In search of an explanation for how a “D/E/Q but not N” N-terminus could have prevented the DsRed2 cargoes from entering the t-ERGIC, we noticed a recent study that reported an analogous rule for protein secretion (Yin et al., 2018): With a growth hormone cargo, it is found that D/E/Q-containing, but not N-containing, N-terminal tripeptides are unfavored for secretion mediated by the receptor SURF4, whereas hydrophobic-proline-hydrophobic (ϕ-P-ϕ) tripeptides are the most favored. Accordingly, we examined an APE-DsRed2-ER-5 variant and found it phenocopied that of EPV-DsRed2-ER-5 (Figures S4A and S4B), thus indicating that the N-terminal tripeptide is also important to cargo sorting into the t-ERGIC. Recent studies on SURF4 and its homologs have generally suggested its preference for hydrophobic N-termini (Belden and Barlowe, 2001; Casler et al., 2019, 2020; Otte and Barlowe, 2004). Therefore, we set out to examine the role of SURF4 in the t-ERGIC pathway.

Live-cell imaging showed that AcGFP1-SURF4 colocalized with the t-ERGICs (Figure 4A, Video S5) and segregated with APV-DsRed2-ER-5 during the *de novo* generation of t-ERGIC (Figure S4C). FLAG-SURF4 also colocalized with the t-ERGIC, and was enriched in the Golgi in APV-DsRed2-ER-5-transfected cells but was mostly in the ER in EPV-DsRed2-ER-5-transfected cells (Figure S4D), suggesting the co-transport of SURF4 and its cargo (Wang et al., 2021). Notably, SURF4-HA co-immunoprecipitated much more efficiently with APV-FLAG-DsRed2-ER-5 than EPV-FLAG-DsRed2-ER-5, even as the input amount of the former was several-fold lower due to secretion (Figure 4B).

**Figure 4.**
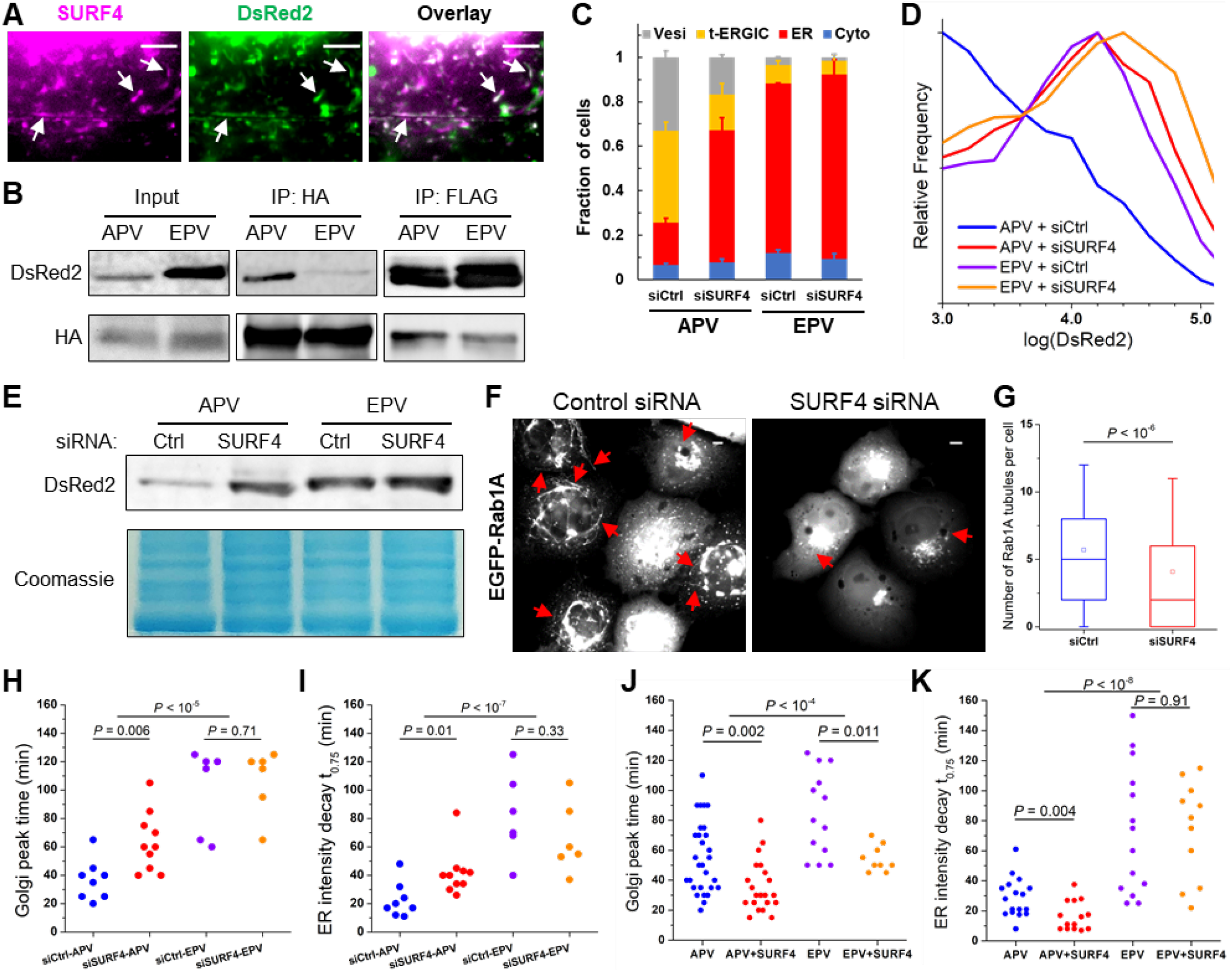
SURF4 Recognizes the N-terminus of the Cargo and Enables t-ERGIC Formation for Expedited ER-to-Golgi Trafficking. (A) Live-cell images of co-transfected AcGFP1-SURF4 and APV-DsRed2-ER-5 in a COS-7 cell. (B) Co-immunoprecipitation of APV/EPV-FLAG-DsRed2-ER-5 with co-expressed SURF4-HA. (C-E) Subcellular distributions (C), flow cytometry histograms (D), and lysate immunoblots (E) of APV/EPV-DsRed2-ER-5 for COS-7 cells co-transfected with control siRNA or SURF4 siRNA. Error bars: SEM (n = 3 with ~50 cells in each replicate). (F,G) Representative fluorescence micrographs (F) and counts per cell (G) of Rab1A-positive tubules in control or SURF4 siRNA-treated COS-7 cells. Whiskers and boxes show 10%, 25%, 50%, 75%, and 90% quantiles. Open squares indicate means. 287 and 321 cells were quantified for siCtrl and siSURF4, respectively. (H,I) Comparison of the time to the peak fluorescence in the Golgi (H) and the time of fluorescence decay to 75% of the start in the ER (I) of APV/EPV-SBP-DsRed2-ER-5 with control siRNA or SURF4 siRNA in RUSH. (J,K) Comparison of the time to the peak fluorescence in the Golgi (J) and the time of fluorescence decay to 75% of the start in the ER (K) of APV/EPV-SBP-DsRed2-ER-5 with or without co-expression of FLAG-SURF4 in RUSH. Scale bars: 5 μm. Arrows point to t-ERGICs. *P* values are calculated by Mann-Whitney test (G), two-way ANOVA (APV vs. EPV in [H-K]), or two-tailed *t* test (siRNA or SURF4 overexpression in [H-K]). See also Figure S4.

With small interfering RNA (siRNA) targeting SURF4 (Figure S4E), we next observed a substantial reduction in APV-DsRed2-ER-5 t-ERGICs (Figures 4C, S4F). The intracellular retention of APV-DsRed2-ER-5, as determined by both flow cytometry and immunoblotting, was also significantly enhanced (Figures 4D and 4E). In comparison, intracellular retention of the SURF4-unfavored EPV variant started high and was only mildly affected by the SURF4 siRNA (Figures 4D and 4E). Notably, in cells not expressing DsRed2 cargoes, SURF4 knockdown also markedly reduced the number of EGFP-Rab1A-labeled t-ERGICs (Figures 4F and 4G). RUSH experiments further showed that SURF4 knockdown substantially decreased the trafficking rate of APV- but not EPV-SBP-DsRed2-ER-5 (Figures 4H and 4I). Conversely, when FLAG-SURF4 was overexpressed, the ER-to-Golgi trafficking of APV-SBP-DsRed2-ER-5 was specifically accelerated (Figures 4J and 4K).

Together, our results indicate that the biogenesis and cargo selectivity of t-ERGIC both depend on SURF4, thus explaining the peculiar “N-terminus rule” we identified for t-ERGIC-based ER-to-Golgi transport.

### Co-clustering of SURF4 and Cargo Expands the ERES for t-ERGIC Biogenesis

To examine how SURF4 facilitated t-ERGIC biogenesis, we utilized STORM to examine whether SURF4 cargoes were sequestered into a special ERES domain. With cells co-transfected with APV-DsRed2-ER-5 and EGFP-Sec23A and immunolabeled for Sec31A, STORM showed that at the ERES (colocalization of Sec23A and Sec31A), Sec31A formed cup-shaped cages (Figures 5A and 5B). Markedly, STORM of APV-DsRed2-ER-5 in a second color channel showed that at the ERES, Sec31A cages that surrounded this SURF4 cargo (e.g., filled arrowheads in Figure 5B) were notably larger than those not loaded with the cargo (e.g., open arrowheads in Figure 5B). As larger ERESs could provide more membrane materials for forming the long t-ERGIC tubules, this observation (statistics in Figure 5C) may explain the specificity of t-ERGIC to SURF4 cargoes. SURF4 siRNA treatment substantially reduced the occurrence of large (>~250 nm) Sec31A cups and hence removed the size difference between DsRed2-loaded and non-loaded ERESs (Figures S5A and S5B), suggesting that SURF4 is necessary for the ERES enlargement in t-ERGIC biogenesis.

**Figure 5.**
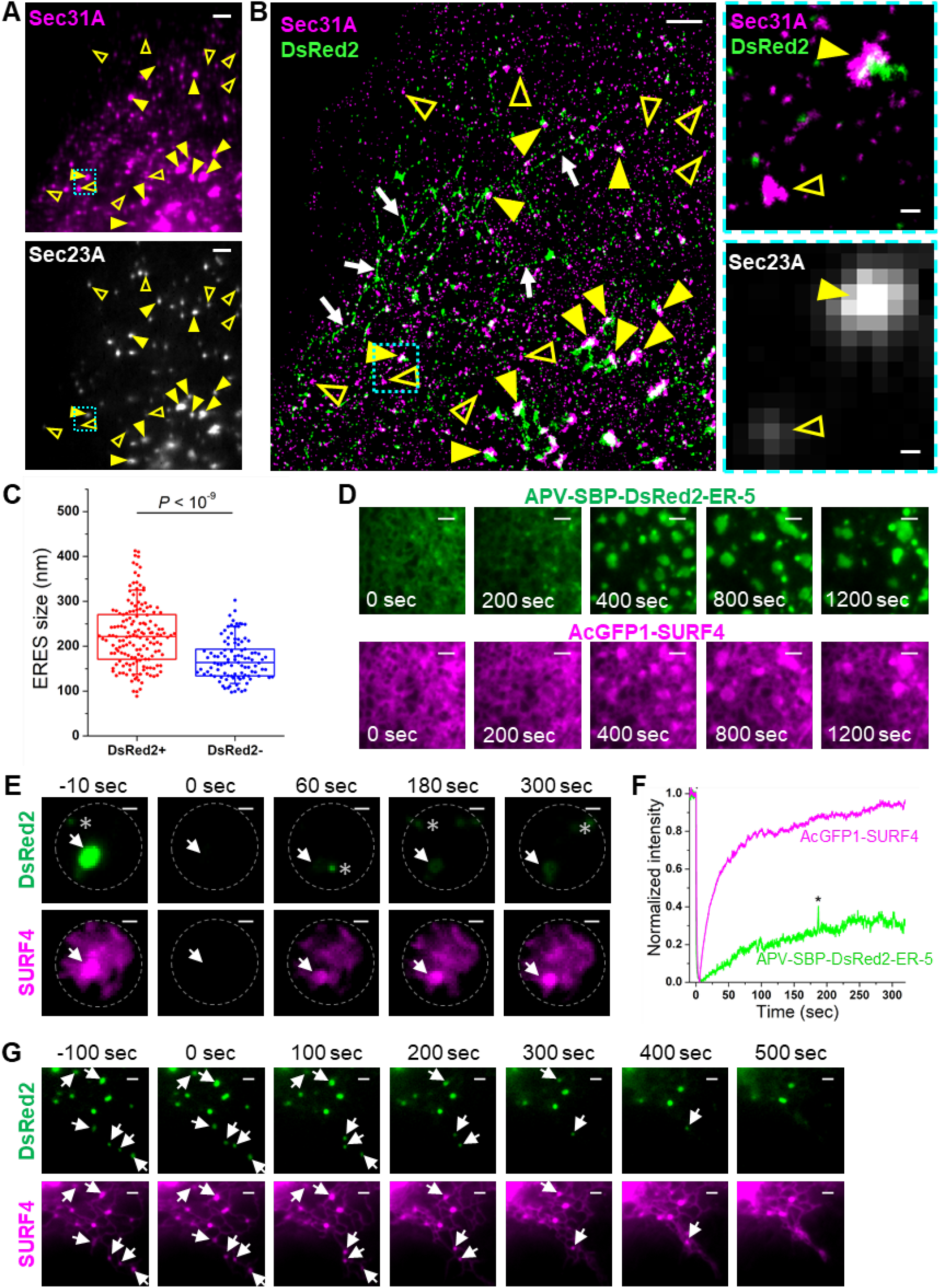
SURF4 Co-clusters with its Cargo to Expand the ERES. (A) Epifluorescence of immunolabeled Sec31A (top) and EGFP-Sec23A (bottom) in a COS-7 cell co-expressing EGFP-Sec23A and APV-DsRed2-ER-5. (B) Two-color STORM image of Sec31A and APV-DsRed2-ER-5 for the same view as (A) (left), as well as zoom-ins (right) of the cyan-boxed region of the STORM image and the EGFP-Sec23A epifluorescence image. Yellow arrowheads in (A,B) indicate examples of ERES labeled with both EGFP-Sec23A and Sec31A. Filled and open arrowheads indicate DsRed2-loaded and non-loaded ERESs, respectively. Arrows in (B) indicate t-ERGICs. (C) Statistics of the sizes of DsRed2-loaded and non-loaded, Sec23A-positive ERESs, based on the STORM-determined sizes of the Sec31A clusters. Whiskers and boxes show 10%, 25%, 50%, 75%, and 90% quantiles. *P* value is calculated by a two-tailed *t* test. n = 5 STORM images were quantified. (D) Representative RUSH image sequence showing the formation and fusion of LLPS-like domains of co-clustered AcGFP1-SURF4 and APV-SBP-DsRed2-ER-5. Biotin was added at time 0 for cargo release. See also Video S7. (E) Dual-color FRAP image sequence of AcGFP1-SURF4 and APV-SBP-DsRed2-ER-5 in a condensate. The gray circle indicates the illuminated area as defined by a pinhole. Asterisks mark random DsRed2-containing vesicles entering the illuminated area. 80 μM biotin was added at −30 min. (F) Fluorescence recovery time trace for the condensate pointed to by the arrow in (E). (G) Dissolution of the AcGFP1-SURF4 and APV-SBP-DsRed2-ER-5 condensates in RUSH by adding 3% 1,6-hexanediol at time 0. Arrows mark the gradually dissolved condensates. 80 μM biotin was added at −30 min. See also Video S8. Scale bars: 2 μm (A,B,D,G), 200 nm (zoom-ins of B), 1 μm (E). See also Figure S5.

To further understand how SURF4 expanded ERES, we turned to live-cell RUSH assay. Upon the release of APV-SBP-DsRed2-ER-5 by biotin, we observed the co-clustering of this cargo with AcGFP1-SURF4 at the ERES, as well as their co-translocation into the fast-moving t-ERGIC (Video S6). However, the transient nature of the cargo-loaded ERES impeded detailed characterization. Interestingly, in a fraction (~20%) of the cells characterized by high expression levels, we observed that after the release of cargo by biotin, SURF4 and the cargo co-clustered to form gradually expanding domains in the ER (Figure 5D, Video S7). 3D-STORM of fixed cells showed that SURF4 and the cargo were both membrane-associated at the expanded clusters (Figures S5C and S5D). These membrane-bound clusters (condensates) showed liquid-like properties as found in liquid-liquid phase separation (LLPS) (Zhao and Zhang, 2020), including their quasi-circular appearance and growth by fusion (Figure 5D; Video S7). After ~15 min, the fusion between the SURF4-cargo condensates slowed down (Video S7), which offered us an opportunity to examine their physical properties in live cells.

We first applied fluorescence recovery after photobleaching (FRAP) to examine the dynamics of SURF4 and cargo in these condensates. Intriguingly, upon photobleaching, whereas the AcGFP1-SURF4 fluorescence quickly recovered in ~1 min, APV-SBP-DsRed2-ER-5 exhibited slow and incomplete recovery (Figures 5E and 5F). This result may be understood as that as SURF4 and its cargo dynamically bound and unbound at the ERES, an excessive amount of the former prevented the diffusion of the latter. We next tested whether the SURF4-cargo condensates could be disrupted by 1,6-hexanediol, an amphiphilic small molecule widely used in LLPS characterizations (Kroschwald et al., 2017). We thus found that as AcGFP1-SURF4 and APV-SBP-DsRed2-ER-5 started to cluster, the addition of 3% 1,6-hexanediol led to gradual, yet always concurrent dissolution of the SURF4 and DsRed2 condensates, so that only a few larger ones persisted after ~400 s (Figure 5G, Video S8). Together, our results suggest that the co-clustering of SURF4 and cargo provides an LLPS-related mechanism to expand the ERES to facilitate t-ERGIC biogenesis.

### Antagonism between SURF4 and KDEL Receptors Regulates the Steady-State Location of Cargo Proteins

While we have elucidated how soluble proteins of different N-termini were differentially selected by SURF4 for entering the t-ERGIC secretion pathway, further experiments indicated another layer of complexity. Specifically, whereas we showed above that APV-DsRed2-ER-5 and APV-GCaMP6s-ER-5 both mainly localized to t-ERGIC tubules at the steady state, analogous constructs of other FPs, including Dendra2, mOrange2, mCherry, EGFP, and mEmerald, mainly localized to the ER and had only ~10% cells dominated by fluorescence in the t-ERGIC (Figures 6A, S6A and S6B).

**Figure 6.**
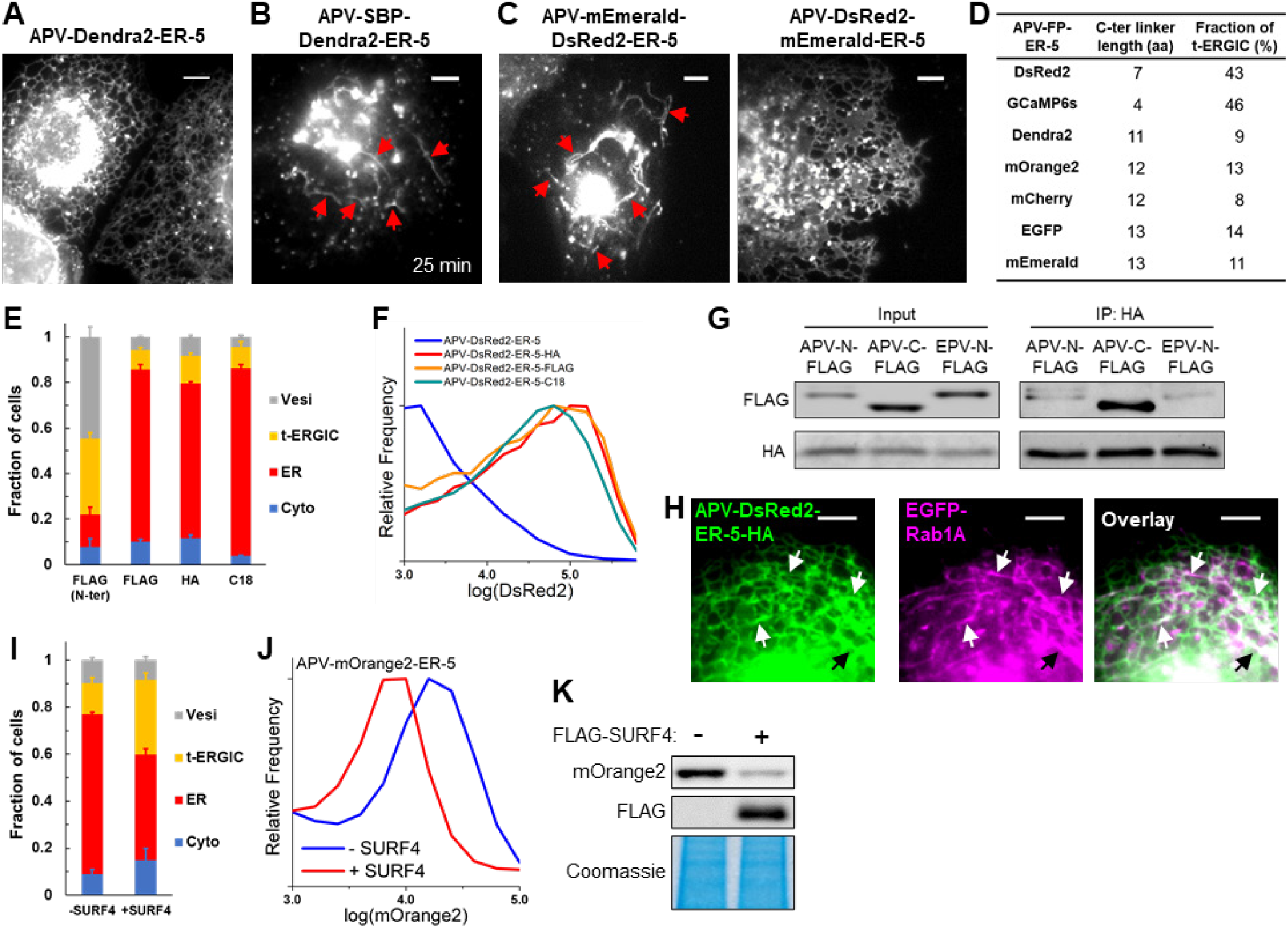
Antagonism between SURF4 and KDELR Determines the Fates of Secretory Cargoes. (A) Representative fluorescence micrograph of APV-Dendra2-ER-5 in COS-7 cells. (B) Representative fluorescence micrograph of APV-SBP-Dendra2-ER-5 in RUSH. 80 μM biotin was added at 0 min. (C) Representative fluorescence micrographs of APV-DsRed2-mEmerald-ER-5 and APV-mEmerald-DsRed2-ER-5 in COS-7 cells. (D) Table summarizing the C-terminal linker lengths and the fractions of t-ERGIC-predominant cells for different APV-FP-ER-5 constructs. See Methods for detail. (E,F) Subcellular distributions (E) and flow cytometry histograms (F) of APV-(FLAG-)DsRed2-ER-5 and its C-terminal inserted derivatives. The “FLAG (N-ter)” data in (E) duplicates “APV” in Figure S3B. (G) Co-immunoprecipitation of APV-FLAG-DsRed2-ER-5, APV-DsRed2-ER-5-FLAG, and EPV-FLAG-DsRed2-ER-5 with KDELR3-HA. (H) Dual-color live-cell fluorescence micrographs of APV-DsRed2-ER-5-HA and EGFP-Rab1A in a co-transfected cell. (I-K) Subcellular distribution (I), flow cytometry histograms (J), and lysate immunoblots (K) of APV-mOrange2-ER-5 in COS-7 cells with or without the co-expression of FLAG-SURF4. Scale bars: 5 μm. Error bars: SEM (n = 3 with ~50 cells in each replicate). Arrows indicate t-ERGICs. See also Figures S6 and S7.

Curiously, RUSH experiments on APV-SBP-FP-ER-5 showed that upon cargo release, all FPs were efficiently trafficked to the Golgi via t-ERGIC (Figures 6B, S6D and S6E). Thus, although all FP cargoes entered the t-ERGIC pathway, other factors modulated their steady-state location. One possibility is that the cargoes were differentially retrieved back from the Golgi to the ER. As the Golgi-to-ER transport is often mediated by KDEL receptors (KDELRs), which recognize KDEL-like motifs at the cargo C-termini (Munro and Pelham, 1987; Wilson et al., 1993), we compared APV-mEmerald-DsRed2-ER-5 and APV-DsRed2-mEmerald-ER-5, which respectively had C-termini identical to that of APV-DsRed2-ER-5 and APV-mEmerald-ER-5 (Figure S6C). Remarkably, at the steady state, we found the former was often in the t-ERGIC (Figure 6C), whereas the latter was mainly in the ER (Figure 6C) and was better retained in the cell (Figure S6C).

To rationalize how the C-termini of APV-DsRed2-ER-5 and APV-mEmerald-ER-5, which both ended with KDEL, could interact differently with KDELRs, we noted that recent structural analysis indicates that the KDELR binding pocket is largely buried inside the membrane (Bräuer et al., 2019). It is thus possible that the binding efficiency of KDELRs may depend on how well the C-terminus KDEL motifs are exposed. Indeed, as we examined the linker between the folded FP core and the C-terminus KDEL motif, we found that the DsRed2 and GCaMP6s constructs had much shorter linkers when compared to the other FPs (Figure 6D, Methods).

To test whether this linker length could be significant, we inserted into APV-DsRed2-ER-5 three different sequences (FLAG-tag of 8 aa, HA-tag of 9 aa, and a random 18-aa linker) between the DsRed2 C-terminus and the KDEL motif (Figure S7A). Remarkably, these constructs of extended pre-KDEL linkers all mainly localized to the ER (Figures 6E, S7B) and showed substantially increased intracellular retention (Figure 6F). Co-immunoprecipitation showed that the C-terminally extended cargo indeed interacted with the KDELR much more strongly when compared to control constructs in which the same extension was added to the N-terminus (Figure 6G). Conversely, as we truncated 10 C-terminal residues before the KDEL motif in APV-EGFP-ER-5, increased t-ERGIC presence of the cargo was observed (Figures S7C and S7D) together with reduced intracellular retention (Figure S7E). Together, our results suggest that the efficacy of KDELR-mediated Golgi-to-ER transport, and hence the steady-state localization of cargoes, depend on how well the C-terminus KDEL motif is exposed.

Notably, although the C-terminally extended APV-DsRed2-ER-5-HA mainly localized to the ER, co-imaging with EGFP-Rab1A showed that it also populated Rab1A-positive t-ERGICs (Figure 6H), whereas EPV-DsRed2-ER-5-HA did not (Figure S7F). Moreover, as we overexpressed FLAG-SURF4 in cells expressing APV-mOrange2-ER-5, more cells were characterized by fluorescence in the t-ERGICs (Figure 6I), and the intracellular retention decreased (Figures 6J and 6K). Thus, the N-terminal SURF4 signal and the C-terminal KDELR signal independently promote anterograde and retrograde trafficking, and thus antagonistically regulate the steady-state localization and retention of the cargo.

## DISCUSSION

Although the molecular diversity of cargo-receptor interactions in the early secretory pathways have been extensively characterized biochemically, their cell biological consequences, including the diversity of cargo carriers and the differential transport kinetics, are less understood (Barlowe and Helenius, 2016; Dancourt and Barlowe, 2010; Gomez-Navarro and Miller, 2016). Our results showed that SURF4-cargo interactions give rise to a morphologically and functionally distinct compartment that specifically expedites the ER-to-Golgi transport of SURF4 cargoes (Figure 7A).

**Figure 7.**
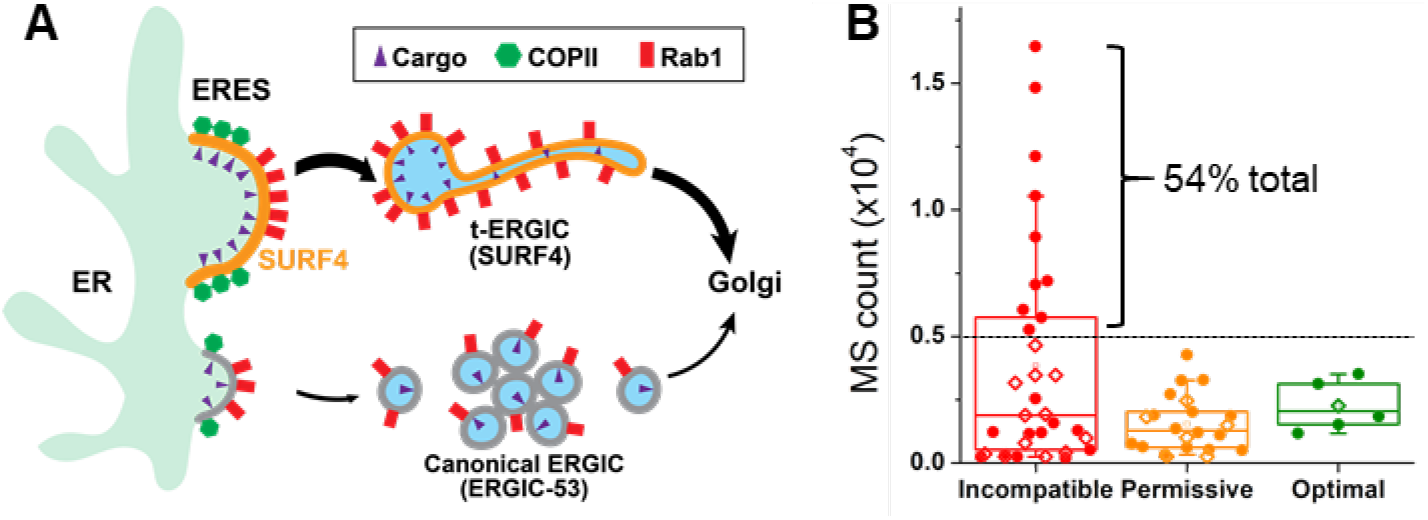
SURF4-Mediated t-ERGIC Transport Leads to Differential Protein Trafficking Rates and Steady-State Localizations. (A) Working model of differential ER-to-Golgi trafficking via the SURF4-mediated t-ERGIC vs. the canonical ERGIC. (B) Categorization of the mass spectrometry counts of ER-lumen proteins in the HeLa cell (Itzhak et al., 2016), based on the N-terminal tripeptide, into SURF4-incompatible (D/E/Q-containing), optimal (ϕ-P-ϕ), and permissive (others). Proteins with and without KDEL-like motifs (Raykhel et al., 2007) are marked by filled circles and open diamonds, respectively.

Whereas studies in yeasts suggest the budding off of COPII-coated vesicles at the ERES as ER-to-Golgi carriers (Dancourt and Barlowe, 2010; Kurokawa and Nakano, 2019; Lee et al., 2004), live imaging of mammalian cells has shown COPII coats stably associate with the ER (Mironov and Beznoussenko, 2019; Shomron et al., 2019; Stephens et al., 2000; Westrate et al., 2020). Our RUSH results are in line with the latter observations. This contrasting behavior may be attributed to the lack of ERGIC in model yeasts, thus highlighting the importance of ERGIC in relaying and sorting cargoes for the much larger mammalian cells.

Early electron microscopy studies depict ERGICs as VTCs with ~100 nm tubular buds extending from vesicular bodies (Saraste and Svensson, 1991; Schweizer et al., 1988). The abundance of ERGIC-53 in the VTCs has since made it a canonical marker for ERGIC (Appenzeller-Herzog and Hauri, 2006; Hauri and Schweizer, 1992; Saraste and Marie, 2018). Although ERGIC-53-positive and ERGIC-53-negative tubular carriers of >2 μm lengths have been observed for certain cargoes (Ben-Tekaya et al., 2005; Blum et al., 2000; Marra et al., 2001; Presley et al., 1997; Sannerud et al., 2006; Shomron et al., 2019; Simpson et al., 2005), it remains unclear what cargo features and/or their molecular interactions lead to this phenotype. Our results showed that the cargo receptor SURF4 defines an ERGIC-53-negative ERGIC domain that is morphologically distinct from VTCs, being ~10 μm long and <30 nm in diameter.

While lacking ERGIC-53, the t-ERGIC is enriched with Rab1. Rab1 plays key roles in ER-to-Golgi trafficking by recruiting motors to enable budding and effectors to mediate targeting and fusion (Allan et al., 2000; Moyer et al., 2001). Our RUSH experiments showed the accumulation and co-budding of Rab1 with the cargo at the ERES for both the *de novo* generation and fusion-elongation of the t-ERGIC, and that a dominant negative Rab1A mutant abolished t-ERGIC generation. These results echo previous findings that Rab1 is indispensable for cargo export (Plutner et al., 1991; Stenmark, 2009; Tisdale et al., 1992), as well as recent experiments reporting the ERES accumulation and co-budding of Rab1 into Golgi-bound carriers (Shomron et al., 2019; Westrate et al., 2020).

The biogenesis of the extraordinarily long t-ERGIC demands a large amount of membrane materials. STORM showed that ERESs loaded by SURF4 cargoes were considerably larger. Moreover, as we depleted SURF4, the enlarged ERESs disappeared and the number of t-ERGIC in the cell diminished. Though a similar drop in ERES size has been noticed with SURF4 knockdown in *C. elegans*, the molecular mechanisms remain unclear (Saegusa et al., 2018). With RUSH live-cell imaging, we captured the gradual expansion of the ERES after cargo release. In particular, for cells with high expression levels, we observed the co-clustering of SURF4 and cargo at large, stable domains characteristic of membrane-bound LLPS condensates. Although the structure of SURF4 is unresolved, recent work has shown that SURF4 oligomerizes *in vivo* (Wang et al., 2021), thus implying possible multivalent interactions required by LLPS (Feng et al., 2019).

Whereas ERES expansion *via* SURF4-cargo interactions provides a potential mechanism for t-ERGIC generation, additional machineries await to be identified to explain the recruitment of Rab1, which we showed to enable both the ERES budding of new t-ERGICs and the fusion with pre-existing t-ERGICs. Packing more cargoes into an existing t-ERGIC is potentially more efficient than transporting many smaller vesicles. The accumulation of membrane materials through the continued fusion with more ERESs also explains how the very long t-ERGICs could form. The extreme lengths and thinness of these carriers may be natural consequences of the Rab1-recruited motors (Stephens, 2012), which rapidly pulled the t-ERGICs along the microtubules.

By systematically comparing the behavior of different cargo proteins under native, SURF4-depleted, and SURF4-overexpressed conditions, we showed that SURF4 selectively routed its cargoes to the t-ERGIC for accelerated ER-to-Golgi transport. Whereas it has been recognized that the SURF4-mediated transport is substantially faster than the bulk flow (Belden and Barlowe, 2001; Casler et al., 2019, 2020; Dancourt and Barlowe, 2010; Emmer et al., 2018; Malkus et al., 2002; Otte and Barlowe, 2004; Saegusa et al., 2018; Yin et al., 2018), our results unveiled that SURF4 establishes a distinct ERGIC form to facilitate this process. By virtue of its *en bloc* cargo packaging, high moving speed, and fast recycling capability, the t-ERGIC provides an efficient trafficking pathway. With its extremely elongated shape and hence high surface-to-volume ratio, the t-ERGIC may be particularly efficient for the transport of receptor-bound cargoes at the membrane while minimizing the nonspecific trafficking of other soluble proteins in the lumen (Saraste and Marie, 2018).

While our RUSH results showed that SURF4 cargoes consistently entered the t-ERGICs for rapid ER-to-Golgi transport, at the steady state some cargoes localized more strongly to the ER, even though Rab1 co-labeling showed that they entered t-ERGICs. Whereas KDELRs provide a well-studied mechanism for retrograde trafficking (Gomez-Navarro and Miller, 2016), we unveiled an interesting effect, in which the C-terminal KDEL motif was less accessed by the KDELRs when closely linked to a well-folded core. Extending this linker substantially increased the KDEL-KDELR affinity, under which condition the cargo became more localized to the ER. Overexpressing SURF4 tipped this balance again and led to more pronounced localization of the cargo in the t-ERGIC at the steady state and decreased intracellular retention. Together, we thus showed that the N-terminus-selective, SURF4-mediated t-ERGIC fast route for ER-to-Golgi transport may be counterbalanced by the C-terminal ER-retrieval signal for regulating the spatiotemporal distribution of the cargo.

For ER-resident soluble proteins, one may thus expect that the SURF4 signal to be negatively selected. We surveyed the N-terminal tripeptides of the ER-resident proteome based on a subcellular fractionation-mass spectrometry dataset of HeLa cells (Itzhak et al., 2016). Notably, out of the 61 annotated ER-lumen proteins, the 10 most abundant ones, making up 54% of the total, all have SURF4-incompatible N-termini together with KDEL-like C-terminal motifs (Figure 7B, Table S1). Low SURF4-binding affinity may thus have been evolutionarily selected for the abundant ER-resident proteins (Yin et al., 2018). Intriguingly, our analysis also identified ER-resident proteins with SURF4-optimal N-termini (Figure 7B, Table S1). Although 5 out of these 6 proteins have KDEL-like ER retrieval motifs (Raykhel et al., 2007), a survey of the literature and our immunofluorescence images both indicated the substantial presence of these proteins outside the ER in the Golgi, vesicles, and the extracellular space (Honoré, 2009; Tsukumo et al., 2009; Vorum et al., 1999). Of note, immunolabeled endogenous calumenin colocalized with EGFP-Rab1A-marked t-ERGIC (Figure S7G), thus suggesting antagonistic trafficking may be utilized by the cell to enrich proteins in the intermediate organelles along secretory pathways.

In summary, by identifying t-ERGIC as a SURF4-mediated, morphologically and functionally distinct compartment that specifically expedites the ER-to-Golgi transport of SURF4 cargoes, our results argue that specific cargo-receptor interactions give rise to distinct transport carriers, which in turn regulate the ER-to-Golgi trafficking kinetics. Given the diversity of cargo receptors, it remains open whether other receptor-cargo interactions may produce yet other ERGIC forms. Meanwhile, the antagonism between the N-terminal ER export and C-terminal ER retrieval signals unveiled in this work demonstrates how the cargo primary structure may be utilized to achieve exquisite, hierarchical controls of protein trafficking and localization.

## ACKNOWLEDGEMENTS

We thank Dr. Mary West of QB3 Cell and Tissue Analysis Facility for technical support with flow cytometry. We acknowledge support by the National Institute of General Medical Sciences of the National Institutes of Health (DP2GM132681). K.X. is a Chan Zuckerberg Biohub investigator and acknowledges additional support by the Packard Fellowships for Science and Engineering.

## AUTHOR CONTRIBUTIONS

R.Y. and K.X. designed the study and wrote the manuscript with input from K.C. R.Y. conducted the experiments and analyzed the data. K.C. contributed to optical setups and initial experiments. K.X. supervised the study.

## ADDITIONAL INFORMATION

Supplementary Information is available for this paper. The authors declare no competing interests.

## METHODS

### Cell Culture

COS-7, U2OS, and HeLa cells were obtained from the Cell Culture Facility at University of California Berkeley. Cells were cultured in Dulbecco’s Modified Eagle Medium (DMEM, Gibco 31053-028) supplemented with 10% fetal bovine serum (FBS, Gibco A3160401), 1x GlutaMax (Gibco 35050061), and 1x non-essential amino acids (Gibco 11140050) at 37°C, 5% CO_2_, and ambient oxygen. Lipofectamine 3000 (Invitrogen L3000008) was used for transient transfection according to the manufacturer’s protocol. In general, cells were plated 20-24 hr before transfection to reach 60%-70% confluency. A total of 1 μg plasmid was used for each sample in a 12-well plate (Corning 3513). Experiments were performed 20-24 hr post-transfection, except for RUSH, where the plasmids were expressed for 40-48 hr to ensure adequate expression.

### Plasmids

The following plasmids were from Addgene: pDendra2-ER-5 (57716), pmEmerald-ER-3 (54082), pAcGFP1-Sec61β (15108), pEGFP-ERGIC-53 (38270), pEGFP-Rab1A (49467), pEGFP-Rab5B (61802), pEGFP-Rab11A (12674), pEGFP-Rab7A (12605), pClover-LAMP1 (56528), pEGFP-p62 (38277), pEGFP-Sec23A (66609), pStr-KDEL_SBP-EGFP-Ecadherin (65286), pcDNA3-FLAG-Rab1A-N124I (46778).

The following plasmids were synthesized by Twist Bioscience: pTwist-CMV BetaGlobin-KDELR3-HA (HA inserted between E143 and A144 of human KDELR3), pTwist-CMV BetaGlobin-SURF4-HA (HA inserted between D263 and K265 of human SURF4).

The Golgi-GFP BacMam construct was from Thermo Fisher (Invitrogen C10592), and was transduced following the manufacturer’s protocol.

pDsRed2-ER-5 was constructed from pDendra2-ER-5 by replacing Dendra2 with DsRed2 using the AgeI and Kpn2I sites. The EPV construct (A18E mutation of APV-DsRed2-ER-5) was generated by changing the alanine codon (GCA) to a glutamate codon (GAA) using the BmtI and AgeI sites of the ER-5 plasmids. N-terminal tags (SBP, FLAG) were inserted using the AgeI site, duplicating the APVAT/EPVAT linker. Other A18X mutations were generated in a similar manner. C-terminal tags (HA, FLAG, C18) were inserted using the Kpn2I site, duplicating the SG linker. Other pFP-ER-5 plasmids were generated by replacing DsRed2 with corresponding FPs. pDsRed2-ER-3 was constructed from pmEmerald-ER-3 by replacing mEmerald-KDEL with DsRed2-KDEL using the AgeI and EcoRI sites. pmCherry-Rab1A was constructed by replacing EGFP of pEGFP-Rab1A by mCherry using the AgeI and Kpn2I sites. pAcGFP1-ERGIC-53 (high expression) and pAcGFP1-SURF4 were constructed by replacing Sec61β of pAcGFP1-Sec61β with ERGIC-53 or SURF4 using the Kpn2I and SalI sites. pStr-KDEL_APV/EPV-SBP-DsRed2-ER-5 was constructed by replacing SBP-EGFP-Ecadherin of pStr-KDEL_SBP-EGFP-Ecadherin with APV/EPV-SBP-DsRed2-ER-5 using the AscI and XbaI sites. pHaloTag-Sec23A was constructed by replacing EGFP of pEGFP-Sec23A with HaloTag using AgeI and Kpn2I sites. pStr-KDEL_TPV-CXCL9-mCherry-SBP was constructed by inserting the synthesized human CXCL9-mCherry-SBP (Twist Bioscience) in between the AscI and XbaI sites. The TPV-to-EPV mutation was generated by PCR between the intrinsic BsrGI site and the XbaI site. pStr-KDEL_APV-Dendra2/mCherry/EGFP-ER-5 was constructed by the Gibson assembly (New England BioLabs E2611) of pStr-KDEL_SBP-EGFP-Ecadherin linearized by AscI and XbaI, PCR-amplified calreticulin signal peptide-SBP tag, and PCR-amplified Dendra2/mCherry/EGFP-SGKDEL. pFLAG-SURF4 was constructed by inserting PCR-amplified FLAG-SURF4 from COS-7 cDNA between the BmtI and EcoRI sites of pDsRed2-ER-5. pAPV-DsRed2-mEmerald-ER-5 and pAPV-mEmerald-DsRed2-ER-5 were made by inserting mEmerald into pDsRed2-ER-5 using the Kpn2I site and the AgeI site, respectively. pAPV-EGFP(1-228)-ER-5 was constructed by replacing the full-length EGFP of pAPV-EGFP-ER-5 with EGFP(1-228) using the AgeI and Kpn2I sites. pAPV/EPV-DsRed2-ER-5-KDELKO were made by removing the KDEL motif in the corresponding plasmids using the Kpn2I and EcoRI sites.

All constructed plasmids were prepared from DH5α, XL1-Blue, or Stbl3 cells (from University of California Berkeley QB3 MacroLab) using the QIAprep Spin Miniprep kit (QIAGEN 27106). Protein-coding sequences were verified by Sanger sequencing at UC Berkeley DNA Sequencing Facility.

### Antibodies

The following secondary antibodies were conjugated in house using previously described protocol (Dempsey et al., 2011): goat anti-mouse (Jackson ImmunoResearch 715-005-151)-CF568 (Biotium 92131), goat anti-mouse IgG2b (Jackson 115-005-207)-Alexa Fluor 647 (A37573), goat anti-mouse IgG1 (Jackson ImmunoResearch 115-005-205)-CF568, goat anti-chick (Jackson ImmunoResearch 703-005-155)-Alexa Fluor 488 (Invitrogen A20000). These antibodies (0.3-0.4 mg/mL) were used at 1:60 dilution for immunofluorescence and 1:300 for immunoblotting.

The following antibodies and dilutions were used for immunofluorescence: rabbit anti-β-COP (Invitrogen PA1-061, 1:100), rabbit anti-Rab1A (Cell Signaling Technology 13075, 1:50), rabbit anti-Rab1B (Proteintech 17824-1-AP, 1:40), mouse anti-DsRed (Santa Cruz 390909, 1:100), rabbit anti-FLAG (Cell Signaling Technology 14793, 1:500), rabbit anti-Sec31A (Proteintech 17913-1-AP, 1:200), mouse anti-calumenin (Santa Cruz 271357, 1:30), rabbit anti-GFP-Alexa Fluor 647 (Invitrogen A31852, 1:300), goat anti-rabbit-Alexa Fluor 647 (Invitrogen A21245, 1:400), goat anti-mouse-Alexa Fluor 647 (Invitrogen A21236, 1:400), goat anti-mouse-CF568 (1:60).

For immunoblotting, the following antibodies and dilutions were used: mouse anti-DsRed (1:300), mouse anti-α-tubulin (Sigma T9026, 1:3000), NeutrAvidin (Thermo Scientific 31000)-Alexa Fluor 647 (1:600), chick anti-α-tubulin (Abcam 89984, 1:600), mouse anti-PERK (Santa Cruz 377400, 1:300), rat anti-GRP94 (Santa Cruz 32249, 1:600), rabbit anti-GRP78 (Invitrogen PA5-29705, 1:1000), rabbit anti-FLAG (1:1000), mouse anti-HA (Invitrogen 26183, 1:2000), rabbit anti-GFP-Alexa Fluor 647 (1:1500), goat anti-rabbit-Alexa Fluor647 (1:2000), goat anti-mouse-CF568 (1:300), goat anti-chick-Alexa Fluor 488 (1:300), goat anti-mouse IgG2b-Alexa Fluor 647 (1:300), goat anti-mouse IgG1-CF568 (1:300).

### Drug Treatments

Cells were transfected for 20-24 hr before the addition of the drug for the indicated time. The following chemicals were used: brefeldin A (Abcam 120299), dithiothreitol (Thermo Scientific R0861), MG132 (Tocris 1748), CB5083 (Cayman Chemical 19311), thapsigargin (Invitrogen T7459), dimethyl sulfoxide (DMSO, Sigma-Aldrich 276855), 1,6-hexanediol (Sigma-Aldrich 240117).

### Live-Cell Fluorescence Microscopy

Cells were plated in Lab-Tek II chambered coverglass (Thermo Scientific 155409) and transfected as described above. For cells transfected with HaloTag-Sec23A, 0.2 μM of JF635 HaloTag ligand (Lavis Lab) was added to the cell culture medium 1 hr before imaging. After incubation at 37°C for 30 min, the cells were rinsed with normal cell culture medium for 5 min × 6 times. Prior to imaging, 25 mM HEPES (Gibco 15630080) was added to the cell culture medium to maintain the pH in the ambient environment.

Live-cell fluorescence microscopy was performed on an Olympus IX73 inverted epifluorescence microscope with a water-immersion objective (Olympus, UPLSAPO60XW, NA 1.2) and a mercury lamp, or a Nikon Eclipse Ti-E inverted fluorescence microscope with an oil-immersion objective (Nikon CFI Plan Apochromat λ 100x, NA 1.45) with 488-nm, 560-nm, and 647-nm lasers modulated by an acousto-optic tunable fiber (AOTF, Gooch & Housego, 97-03151-01). Cells were imaged at 2-20 frames per second (fps) at room temperature to moderately slow down the motion of the fast-moving t-ERGIC. Concurrent multi-color imaging was achieved by modulating the AOTF to allow frame-synchronized alternating excitation at 488, 560, and 647 nm (Yan et al., 2020) with a multi-bandpass filter cube (Semrock Di01-R405/488/561/635 and Chroma ZET405/488/561/640m).

### Single-Particle Tracking Analysis

The time-sorted image sequence was imported into Fiji (Schindelin et al., 2012) and analyzed through the TrackMate plugin (Tinevez et al., 2017) with the LoG detector and the simple LAP tracker. For t-ERGICs, the vesicular bodies were tracked as single particles, and the tubules were not separately tracked. The tracked trajectories were outputted to MATLAB for plotting into scaled displacements of trajectories. The Golgi territory was manually defined.

### RUSH Assay and Analysis

Constructs for the RUSH assay (pStr-KDEL plasmids with a co-expressed streptavidin-KDEL hook) were modified from previous work (Boncompain et al., 2012). Cells were plated in 8-well Lab-Tek chambered coverglass and transfected. The cells were imaged in the cell culture medium with 25 mM HEPES on the abovementioned Nikon Ti-E microscope. 80 μM D-biotin (J&K Scientific 322564) was added to the imaging medium to release the cargo. For quantification of the ER-to-Golgi transport kinetics, images were acquired every 5 min for ~10 predetermined positions using Micro-Manager. For high temporal resolution imaging, images were acquired continuously at 2-20 fps.

To analyze the ER-to-Golgi trafficking rate, the Golgi area and the ER area were manually defined in the image before biotin addition using Fiji. The background-subtracted intensities in the two regions were plotted as a function of time and normalized to the initial values. Golgi peak time was defined as the time corresponding to the highest intensity in the Golgi region. ER intensity decay t_0.75_ was defined as the time when the ER intensity dropped to 0.75 of the initial value. For the very slow ER intensity decay of EPV-DsRed2-ER-5 samples, extrapolation was used to estimate the t_0.75_.

### Cell Fixation, Immunolabeling, and Epifluorescence Microscopy

Cells were plated in Lab-Tek II chambered coverglass or on 12-mm #1.5 coverslips in 24-well plates (Corning 3526) and transfected as described above. Cells were fixed with 3% paraformaldehyde (Electron Microscopy Sciences 15714) with 0.02%-0.1% glutaraldehyde (GA, Electron Microscopy Sciences 16720) in DPBS (Corning 21-030-CV) for 30 min at room temperature. We found that the t-ERGIC morphology was best preserved in the presence of GA, yet a high concentration of GA impeded epitope immunolabeling, especially for antibodies against β-COP and Rab1A. The sample was then reduced with 0.1% NaBH_4_ (Sigma-Aldrich 213462) in DPBS for 5 min, and rinsed with DPBS for 10 min × 3 times.

For immunolabeling, the cells were blocked with the blocking buffer (3% bovine serum albumin [BSA, Sigma-Aldrich A3059] and 0.1% saponin [Sigma-Aldrich S4521] dissolved in DPBS) for 1 hr at room temperature. Primary antibodies were diluted in the blocking buffer at the abovementioned ratios. Cells were incubated with the primary antibodies for 1 hr at room temperature or overnight at 4°C. Cells were then rinsed with the washing buffer (0.1x blocking buffer diluted in DPBS) for 10 min × 3 times before incubation with the secondary antibodies diluted in the blocking buffer for 1 hr at room temperature. After the secondary labeling, cells were rinsed with the washing buffer for 10 min × 3 times and finally with DPBS for 10 min.

Conventional epifluorescence microscopy was performed in DPBS on the same setups for live-cell fluorescence microscopy.

### STORM Super-Resolution Microscopy

STORM experiments were conducted as previously described (Gorur et al., 2017; Hauser et al., 2018). Briefly, the sample was immersed in a photoswitching buffer (5% D-(+)-glucose [Sigma-Aldrich G7528], 100 mM cysteamine [TCI A0648], 0.8 mg/mL glucose oxidase [Sigma-Aldrich G2133], and 40 μg/mL catalase [Sigma-Aldrich C30] in 100 mM Tris-HCl pH 7.5 [Corning 46-030-CM]) and mounted on a custom-built STORM microscope with a cylindrical lens for 3D-STORM (Huang et al., 2008). Single-molecule images were collected at 110 fps for 50,000-80,000 frames for the construction of each super-resolution image. For dual-color STORM, Alexa Fluor 647 and CF568 were sequentially imaged as described above with the 647-nm laser and the 560-nm laser, respectively.

### ERES Size Measurement

STORM single-molecule coordinates of each Sec31A cluster were plotted in the imaging plane, whose long and short axes were defined by a principal direction algorithm as described (Yan et al., 2020). Distributions along the long axis and the short axis were respectively fitted by Gaussian curves. The average of the FWHMs of the two Gaussians was taken as the estimated size of the ERES.

### Flow Cytometry

Flow cytometry was carried out on an Attune NxT flow cytometer (Thermo Fisher) per the manufacturer’s protocols. Transfected cells were trypsinized, neutralized with DMEM, and transferred to a 1.7 mL microcentrifuge tube. ~50,000 cells were analyzed. At 30%-60% transfection efficiency, non-transfected cells in the sample were thresholded by the fluorescence signal and not shown in the graphs.

### Protein Gel Electrophoresis, Coomassie Blue Staining, and Immunoblotting

Cells were plated and transfected in 12-well plates. 20-24 hr after transfection, the cell culture medium was centrifuged at 2,000 g for 2 min and collected for the analysis of secreted proteins (see Immunoprecipitation [IP] and Co-IP below). Cells on the surface of the plate were then lysed in the Triton lysis buffer (1% TritonX-100 [Sigma-Aldrich T8787], 137 mM NaCl [Sigma-Aldrich S9888], 50 mM Tris-HCl pH 7.5, 1x protease inhibitor cocktail [Thermo Scientific 87786], and 1x phosphatase inhibitor cocktail [Sigma-Aldrich P0044] in water) for 30 min on ice. The lysate was then centrifuged at 16,000 g for 15 min at 4°C. The supernatant was added to 1x LDS sample buffer (Invitrogen NP0007) with 300 mM DTT and incubated for 10 min at 75°C.

Samples were run in NuPAGE Bis-Tris gels (4%-12% [Invitrogen NP0321] for small volumes of samples, 10% [Invitrogen NP0315] for larger amounts) in 1x MOPS SDS running buffer (Invitrogen NP0001) at 90 V for 1-2 hr.

For Coomassie Blue staining, the gel was rinsed in water for 5 min, incubated with 50 mL Bio-Safe Coomassie Blue Stain (Bio-Rad 1610786) for 2 hr, and rinsed with water for 30 min × 3 times.

For immunoblotting, the sample in the gel was transferred to a low-fluorescence PVDF membrane (Thermo Scientific 22860) in the transfer buffer (25 mM Tris base [Acros Organics 42457-1000], 192 mM glycine [Sigma-Aldrich G8898], and 10% v/v methanol [VWR BDH1135] in water) at 18 V for 50-70 min at room temperature using the Mini Gel system (Invitrogen NW2000). The membrane was blocked in the blocking buffer of 5% BSA in TBST (137 mM NaCl, 2.7 mM KCl [Sigma-Aldrich P9541], 19 mM Tris-HCl pH 7.5, and 0.1% v/v Tween 20 [Sigma-Aldrich P7949] in water) for 1 hr at room temperature. Primary and dye-conjugated secondary antibodies were diluted in the blocking buffer and incubated with the membrane for 1 hr each. After each round of labeling, the membrane was washed in TBST for 10 min × 3 times. A laboratory rocker (Bellco Biotechnology 7740-10010) was used for all steps. The fluorescently labeled membrane was imaged by the Typhoon FLA 9500 scanner (GE Healthcare Life Sciences) per the manufacturer’s protocol.

### Pulse-Chase Assay

Azidohomoalanine-based pulse-chase assay was performed according to a previous protocol (Wang et al., 2017). Cells were plated and transfected in 6-cm dishes (Falcon 353004). Pulse-chase was performed by incubating the cells in methionine/cysteine/glutamine-free DMEM (Gibco 21013024) with 10% dialyzed FBS (Gibco A3382001), 1x GlutaMax, 0.2 mM L-cysteine (Alfa Aesar J63745), and 50 μM L-azidohomoalanine (AHA, Click Chemistry Tools 1066) for 1 hr at 37°C. The medium was then replaced by the normal cell culture medium supplemented with 2 mM L-methionine (Alfa Aesar J61904) for 0-3 hr at 37°C for the chase. The cells were then lysed in the sodium dodecyl sulfate (SDS) lysis buffer (1% SDS [Sigma L6026], 100 mM Tris-HCl pH 8.0 [Corning 46-031-CM] and 1x protease inhibitor cocktail in water) for 30 min at 4°C, and then centrifuged at 16,000 g for 15 min at 4°C.

Labeling of the incorporated AHA by biotin-alkyne (Click Chemistry Tools 1266) was conducted using the Click-&-Go Protein Reaction Buffer Kit (Click Chemistry Tools 1262) according to the manufacturer’s protocol. The reacted mixture was dialyzed in PBST buffer (DPBS with 0.05% v/v Tween 20) by a centrifugal filter with a 3 KDa molecular weight cut-off (Millipore UFC500324). The supernatant was immunoprecipitated by custom-made anti-RFP-beads (see Immunoprecipitation [IP] and Co-IP below). Biotinylated proteins were detected by western blotting using NeutrAvidin-Alexa Fluor 647.

### Immunoprecipitation (IP) and Co-IP

IP was used to enrich target proteins from the cell culture medium and pulse-chased samples. For anti-FLAG IP of the secreted APV/EPV-FLAG-DsRed2-ER-5, 10-20 μL rat anti-FLAG magnetic beads (Thermo Scientific A36797) were added to 1 mL of the centrifuged cell culture medium (above). For anti-DsRed IP of the AHA pulse-chased samples (above), rabbit anti-RFP magnetic beads were prepared by conjugating 5 μg rabbit anti-RFP antibody (Rockland 600-401-379) to 50 μL Protein G magnetic beads (Invitrogen 10003D) according to the manufacturer’s protocol, and 20-50 μL conjugated beads was used for IP.

Co-IP was performed for cell lysates from 10-cm dishes (Corning 430167). Cells were lysed in the digitonin lysis buffer (1% digitonin [Sigma-Aldrich D141], 137 mM NaCl, 10% w/v glycerol [Alfa Aesar 38988], and 1x protease inhibitor cocktail in water) with pH controlled at 7.0 (for co-IP of APV/EPV-FLAG-DsRed2-ER-5 and SURF4-HA, 17 mM Na_2_HPO_4_ [Macron Fine Chemicals 7917-04], 13 mM NaH_2_PO_4_ [Fisher Chemical S369]) or 6.5 (for co-IP with KDELR3-HA, 18 mM Na_2_HPO_4_, 32 mM NaH_2_PO_4_) for 30 min at 4°C. The lysate was centrifuged at 16,000 g for 20 min at 4°C, and the supernatant was diluted 2x with 200 mM NaCl with 1x protease inhibitor cocktail for co-IP. The rat anti-FLAG magnetic beads and the mouse anti-HA magnetic beads (Thermo Scientific 88836) were used for co-IP.

The sample-loaded beads in 1.7 mL tubes were incubated on a tube rotator (VWR 10136-084) for 3 hr at 4°C. The beads were then washed in the washing buffer (corresponding lysis buffer without digitonin and glycerol supplemented with 0.03% v/v Tween 20) on ice for 5 min × 3 times. The IP-ed proteins were eluted by 20 μL 1.5x LDS sample buffer diluted in the washing buffer for 10 min × 2 times at 80°C, with intermittent vortex mixing.

### RNA Interference

Silencer Select siRNA against SURF4 was purchased from Thermo Fisher Scientific (Ambion 4427037-s13651). Scrambled Silencer Select control siRNA (GUACCAAUUCGUAAGUGUUTT; AACACUUACGAAUUGGUACTT) was synthesized by Thermo Fisher Scientific. Cells were plated in 6-well plates (Corning 3516). siRNA transfection was conducted using Lipofectamine RNAiMAX (Invitrogen 13778) per manufacturer’s protocol. Cells were replated on day 3 to ~70% confluency and transfected with plasmids using Lipofectamine 3000 on day 4.

### RT-PCR Assay

siRNA-transfected cells were harvested on Day 5 by trypsinization. Total RNA was extracted with the RNeasy Mini kit (QIAGEN 74104). Reverse transcription was performed using the GoScript Reverse Transcription Kit (Promega A50001) per manufacturer’s protocol. 20 ng cDNA was used for PCR amplification with the iProof High-Fidelity PCR Kit (Bio-Rad 1725330) for 24 cycles. The PCR product was analyzed in 1.5% agarose (Lonza 50002) gel stained by SYBR Safe (Invitrogen S33102) and imaged by the Typhoon TLA 9500 scanner.

Primers used for PCR were: SURF4 (5’: CTGCTCCTAGCAGAATCCC; 3’: TGCATGGGCTTGTAGACTG); GAPDH (5’: CATCACCATCTTCCAGGAGC; 3’: GGATGATGTTCTGGAGAGCC).

### Dual-Color FRAP and Analysis

FRAP experiments were performed on the Nikon Ti-E setup with wide-field illumination and recording. DsRed2 signal and AcGFP1 signal were simultaneously acquired with frame-synchronized alternating excitation at 560 and 488 nm using the abovementioned AOTF-controlled illumination scheme (see Live-Cell Fluorescence Microscopy above) at 10 fps. Identified region of interest was moved to the center of the view, and an adjustable aperture in the incident light path was closed down into a pinhole to limit illumination to a circular region of ~5 μm diameter. Photobleaching was achieved by setting the intensities of both the 560 and 488 nm laser to ~2 kW/cm^2^ for 5-10 sec. The excitation intensities were then lowered to ~1 W/cm^2^ to record fluorescence recovery. Intensity profiles at the phase-separated condensates were background-subtracted and analyzed in Fiji and replotted by Origin 8.5 (OriginLab).

### Estimation of C-Terminal Linker Lengths of FPs

Crystal structures of relevant FPs were identified in the Protein Data Bank (PDB). We defined the free C-terminal linker as the C-terminal tail extending from the folded core of the protein in the structure, plus the unresolved sequence of the FP and the “SG” linker in the ER-5 constructs. The number of amino acid residues was counted as the length of the C-terminal linker. PDB structures used are: 1G7K (DsRed, for DsRed2), 3WLC (GCaMP6m, for GCaMP6s), 2VZX (Dendra2, for Dendra2), 2H5Q (mCherry, for mOrange2 and mCherry), and 2Y0G (EGFP, for EGFP and mEmerald).

### Quantification and Statistical Analysis

The sample size was not predetermined by statistical methods. Fluorescence images and immunoblots were representative of at least three biological replicates. The sample size and significance test of statistical analyses were indicated in the figure legends. The value of “n” corresponds to the number of biological replicates. A *P* value lower than 0.05 was considered statistically significant.

## SUPPLEMENTAL FIGURES

**Figure S1.**
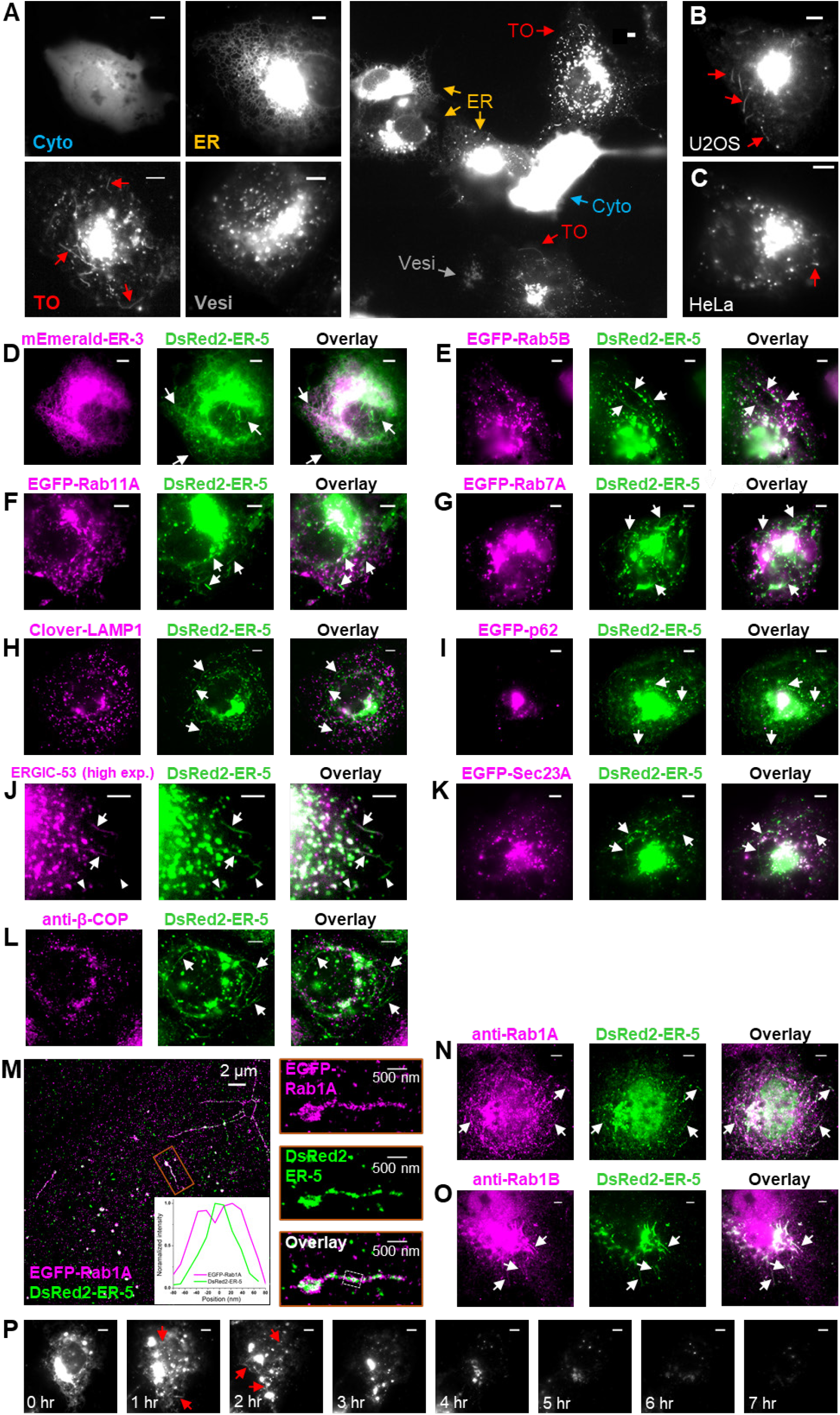
Characterization of DsRed2-ER-5-Containing Tubular Organelles, Related to Figure 1. (A) Representative images of the four types of subcellular distribution of DsRed2-ER-5 in COS-7 cells: cytoplasm (Cyto), ER, tubular organelles (TOs), and vesicles (Vesi). (B,C) Representative images of DsRed2-ER-5 TOs in U2OS (B) and HeLa (C) cells. (D-K) Dual-color fluorescence micrographs of DsRed2-ER-5 (green) in live COS-7 cells with the ER markers mEmerald-ER-3 (D), the early endosome marker EGFP-Rab5B (E), the recycling endosome marker EGFP-Rab11A (F), the late endosome marker EGFP-Rab7A (G), the lysosome marker Clover-LAMP1 (H), the autophagosome marker EGFP-p62 (I), the highly expressed ERGIC marker AcGFP1-ERGIC-53 (J), and the ERES marker EGFP-Sec23A (K). Arrowheads in (J) indicate TOs negative for ERGIC-53. (L) Immunofluorescence of the COPI vesicle marker β-COP vs. DsRed2-ER-5. (M) Two-color STORM of immunolabeled DsRed2-ER-5 and EGFP-Rab1A showing that Rab1A decorates the surface of the TO. Inset: cross-sectional intensity profiles of the boxed region in the overlay. (N,O) Immunofluorescence of endogenous Rab1A (N) and Rab1B (O) in COS-7 cells, showing good colocalization with the DsRed2-ER-5 TOs. (P) Time-lapse imaging of DsRed2-ER-5 in an unsynchronized COS-7 cell, showing its redistribution from the ER to the TOs, Golgi, and vesicles accompanied by a reduction of fluorescence intensity. Scale bars: 5 μm except for the labeled scale bars in (M). Arrows point to TOs.

**Figure S2.**
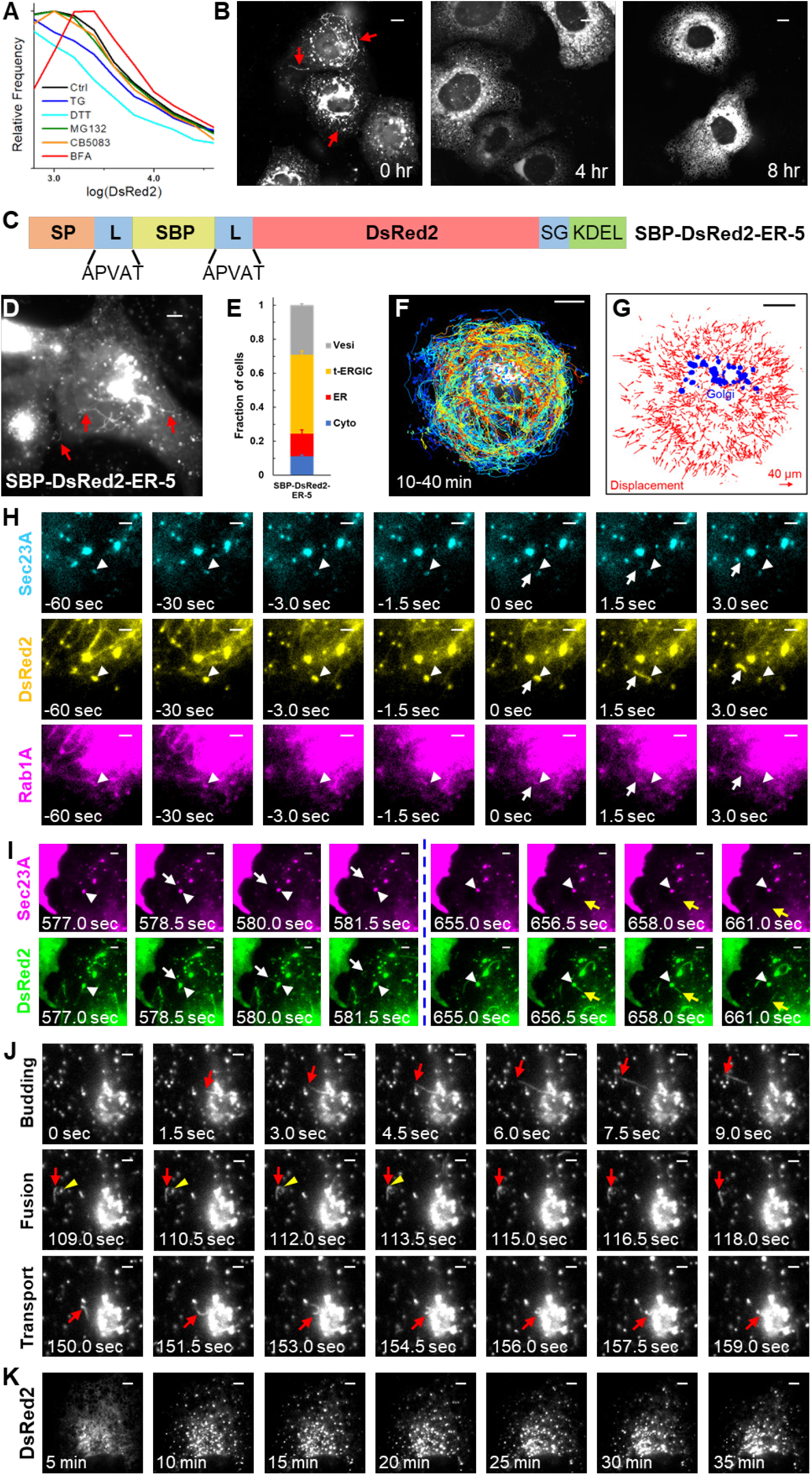
The t-ERGIC Mediates ER-to-Golgi Trafficking by Shuttling between the ER and the Golgi, Related to Figure 2. (A) Flow cytometry histograms of DsRed2-ER-5-transfected COS-7 cells treated with 0.1% DMSO (Ctrl), 1 μM thapsigargin (TG), 7 mM dithiothreitol (DTT), 5 μM MG132, 1 μM CB-5083, or 1 μM brefeldin A (BFA) for 4 hr. (B) Representative fluorescence micrographs of DsRed2-ER-5 in COS-7 cells with brefeldin A treatments of 0, 4, and 8 hr. (C-E) Schematic of SBP-DsRed2-ER-5 (C), a representative image of its presence in the t-ERGIC (D), and its subcellular distribution (E) in transfected COS-7 cells. Error bars: SEM (n = 3 with ~50 cells in each replicate). (F) Single-particle tracking of post-ER carriers in Figure 2D. The color of each trajectory encodes the maximum speed reached. (G) Displacement map of the trajectories in (F). The Golgi apparatus is marked blue. Arrows point to the direction of displacement (from the initial position to the final position), and their magnitudes are scaled according to the legend. (H) Another example of *de novo* formation of SBP-DsRed2-ER-5 t-ERGIC (arrow) from the ERES (arrowhead) in a RUSH experiment, similar to Figure 2F. 80 μM biotin was added at time −11 min for cargo release. (I) RUSH image sequence showing that the same COPII-coated ERES (arrowhead) sequentially generates two t-ERGICs (white and yellow arrows) in opposite directions. 80 μM biotin was added at time 0. (J) RUSH image sequence showing that a t-ERGIC (arrow) buds from the Golgi apparatus, fuses with an ERES (arrowhead), and carries the cargo back to the Golgi apparatus. 80 μM biotin was added 30 min before time 0. (K) Co-transfection of SBP-DsRed2-ER-5 with a dominant negative Rab1A (Rab1A-N124I) inhibited the generation of t-ERGIC in RUSH, and the cargo was stuck at the ERES. 80 μM biotin was added at time 0. Scale bars: 10 μm (B,F,G); 5 μm (D,K); 2 μm (H-J). Arrows in (B,D) indicate t-ERGICs.

**Figure S3.**
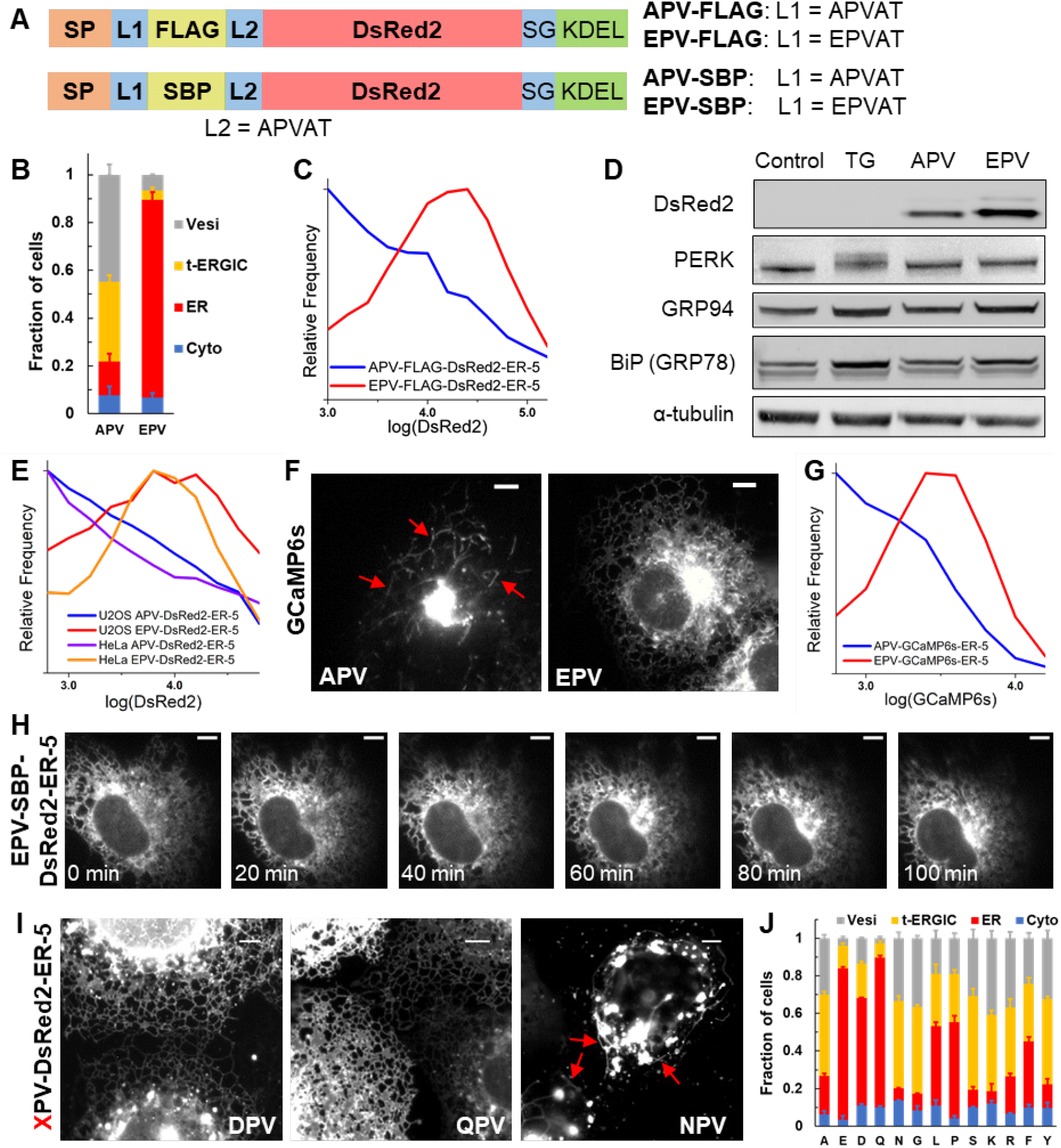
The N-terminus Rule of ER-to-Golgi Transport by t-ERGIC Applies to Different Cargoes, Related to Figure 3. (A) Sequences of APV/EPV-FLAG-DsRed2-ER-5 and APV/EPV-SBP-DsRed2-ER-5. (B,C) Subcellular distributions (B) and flow cytometry histograms (C) of APV/EPV-FLAG-DsRed2-ER-5. (D) Immunoblots of APV/EPV-DsRed2-ER-5-transfected cells and non-transfected cells with or without 1 μM thapsigargin (positive control for ER stress) treatment for 5 hr. (E) Flow cytometry histograms of APV/EPV-DsRed2-ER-5 in HeLa and U2OS cells. (F,G) Representative fluorescence micrographs (F) and flow cytometry histograms (G) of APV/EPV-GCaMP6s-ER-5 in COS-7 cells. (H) Representative RUSH image sequence of EPV-SBP-DsRed2-ER-5. 80 μM biotin was added at time 0. (I) Representative fluorescence micrographs of DPV/QPV/NPV-DsRed2-ER-5 in COS-7 cells. (J) Subcellular distributions of different XPV-DsRed2-ER-5 variants. The “A” and “E” data duplicates that of “ER-5” in Figure 1D and that of Figure 3C, respectively. Scale bars: 5 μm. Error bars: SEM (n = 3 with ~50 cells in each replicate). Arrows point to t-ERGICs.

**Figure S4.**
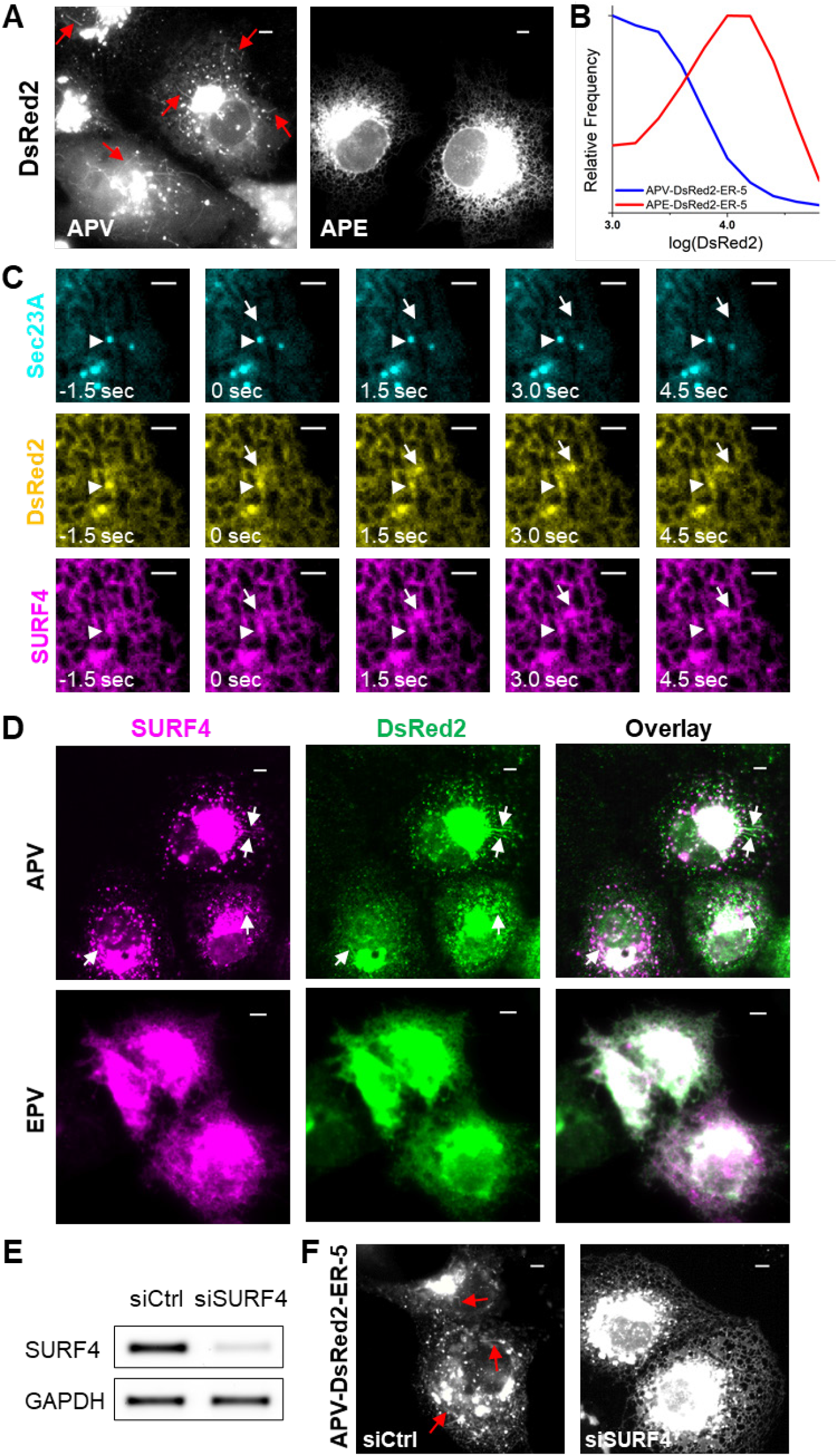
SURF4 Recognizes the N-terminus of the Cargo and Co-traffics with its Selected Cargo via t-ERGICs, Related to Figure 4. (A,B) Representative fluorescence micrographs (A) and flow cytometry histograms (B) of APV/APE-DsRed2-ER-5 in COS-7 cells. (C) Image sequence showing *de novo* generation of a t-ERGIC through the co-budding of AcGFP1-SURF4 (magenta) and APV-DsRed2-ER-5 (yellow) but not JF635-labeled HaloTag-Sec23A (cyan). Arrowhead marks the ERES. Arrow indicates the t-ERGIC. (D) Representative immunofluorescence images of FLAG-SURF4 (magenta) in COS-7 cells co-transfected with APV/EPV-DsRed2-ER-5 (green). (E) RT-PCR of SURF4 mRNA in control and SURF4 siRNA-treated cells. (F) Representative live-cell images of APV-DsRed2-ER-5 in control and SURF4 siRNA-treated cells. Arrows indicate t-ERGICs. Scale bars: 5 μm (A,D,F); 2 μm (C).

**Figure S5.**
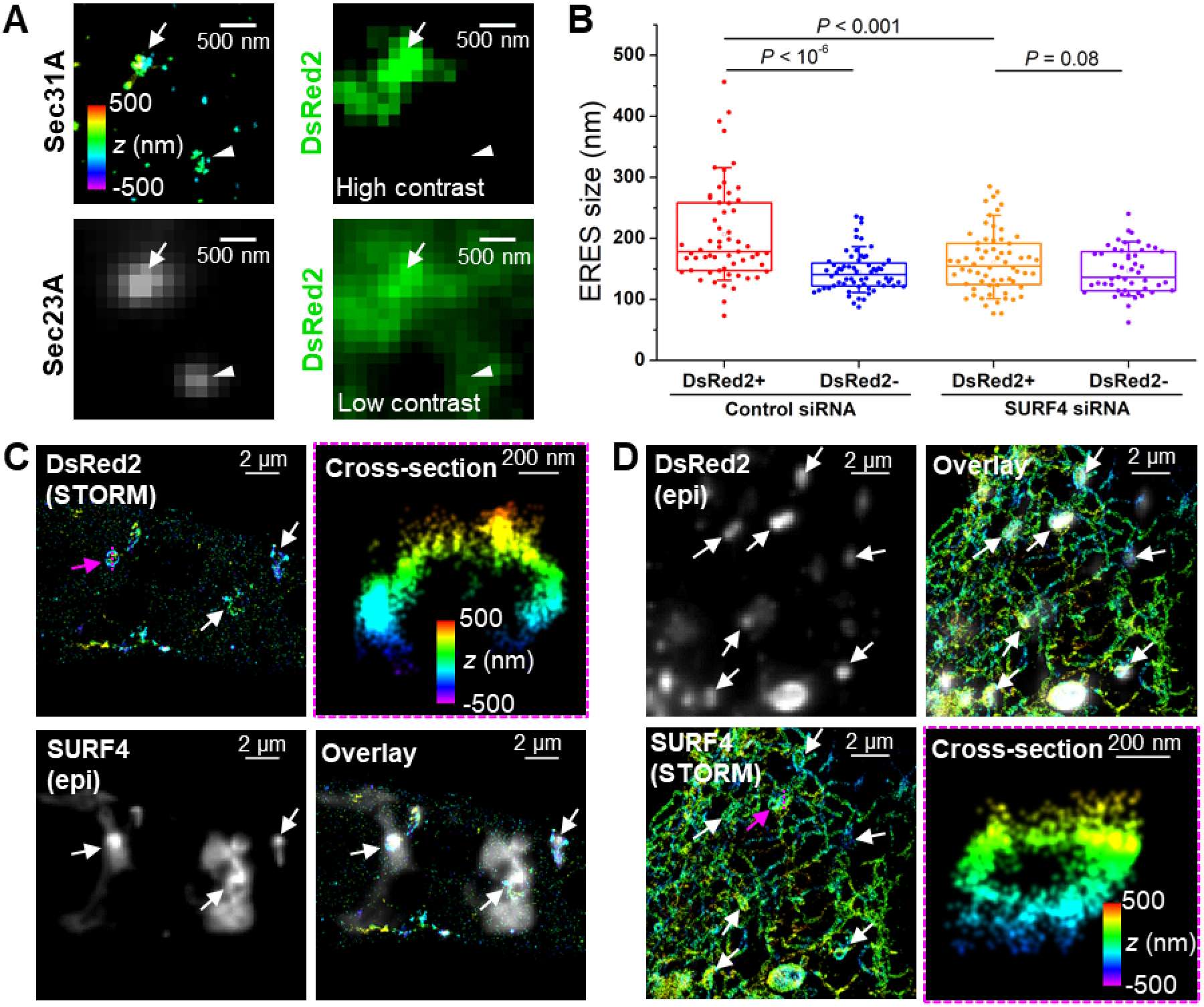
SURF4 and its Cargo Co-cluster to Expand the ERES, Related to Figure 5. (A) DsRed2-loaded (arrow) and non-loaded (arrowhead) ERESs in a SURF4 siRNA-treated cell. Sec31A is shown in 3D-STORM, and epifluorescence of EGFP-Sec23A and APV-DsRed2-ER-5 are shown in gray and green, respectively. Note that with SURF4 knockdown, the APV-DsRed2-ER-5 cargo no longer clustered strongly at the ERES, so that DsRed2-loaded and non-loaded ERESs were subjectively assigned based on enhanced contrast of the epifluorescence image. (B) Statistics of the sizes of DsRed2-loaded and non-loaded, Sec23A-positive ERESs, based on the STORM-determined sizes of the Sec31A clusters, in control and SURF4 siRNA-treated cells. Whiskers and boxes show 10%, 25%, 50%, 75%, and 90% quantiles. *P* values are calculated by the two-tailed *t* test. n = 4 STORM images were quantified. (C,D) 3D-STORM of immunolabeled APV-SBP-DsRed2-ER-5 (C) or AcGFP1-SURF4 (D), in comparison with epifluorescence images of AcGFP1-SURF4 (C) or APV-SBP-DsRed2-ER-5 (D) of the same views in a RUSH experiment. For cargo release, 80 μM biotin was added 40 min before sample fixation. Arrows indicate SURF4 and DsRed2 condensates. Vertical cross-sections of the STORM images are given for the condensates indicated by the magenta arrows, showing membrane localizations for both proteins. Colors in the 3D-STORM images encode axial positions (depth).

**Figure S6.**
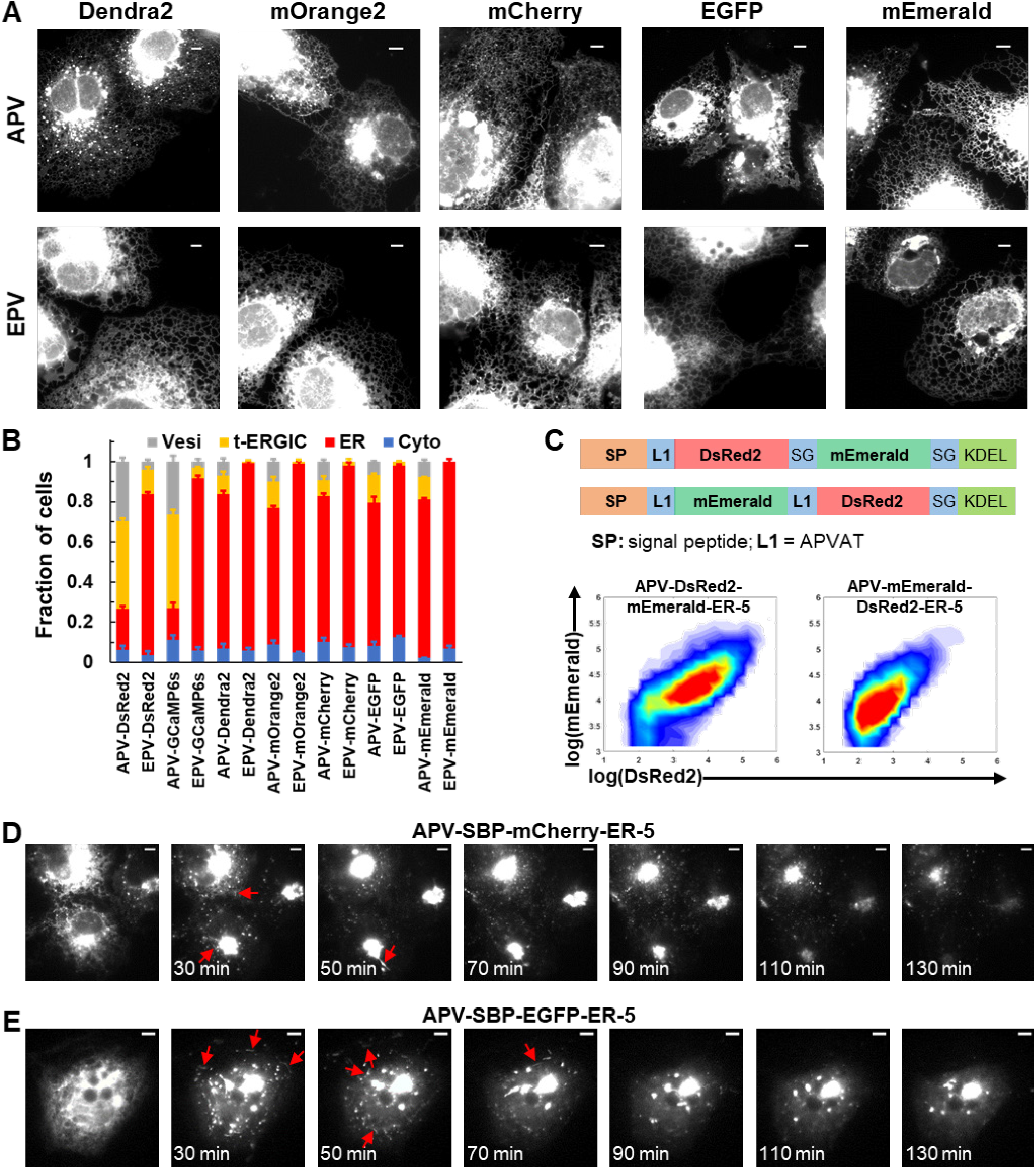
Distinct Steady-State Localizations of Different FP-ER-5 Constructs, Related to Figure 6. (A) Representative fluorescence micrographs of APV/EPV-Dendra2-ER-5, APV/EPV-mOrange2-ER-5, APV/EPV-mCherry-ER-5, APV/EPV-EGFP-ER-5, and APV/EPV-mEmerald-ER-5 in COS-7 cells. (B) Subcellular distributions of different APV/EPV-FP-ER-5 constructs. The APV-DsRed2 and EPV-DsRed2 data duplicate that of “ER-5” in Figure 1D and that of Figure 3C, respectively. The APV-mOrange2 data duplicates that in Figure 6I. Error bars: SEM (n = 3 with ~50 cells in each replicate). (C) Sequences and flow cytometry of APV-DsRed2-mEmerald-ER-5 and APV-mEmerald-DsRed2-ER-5. (D,E) RUSH image sequences of APV-SBP-mCherry-ER-5 (D) and APV-SBP-EGFP-ER-5 (E) showing efficient ER exit and the formation of t-ERGIC. Arrows indicate t-ERGICs. Scale bars: 5 μm.

**Figure S7.**
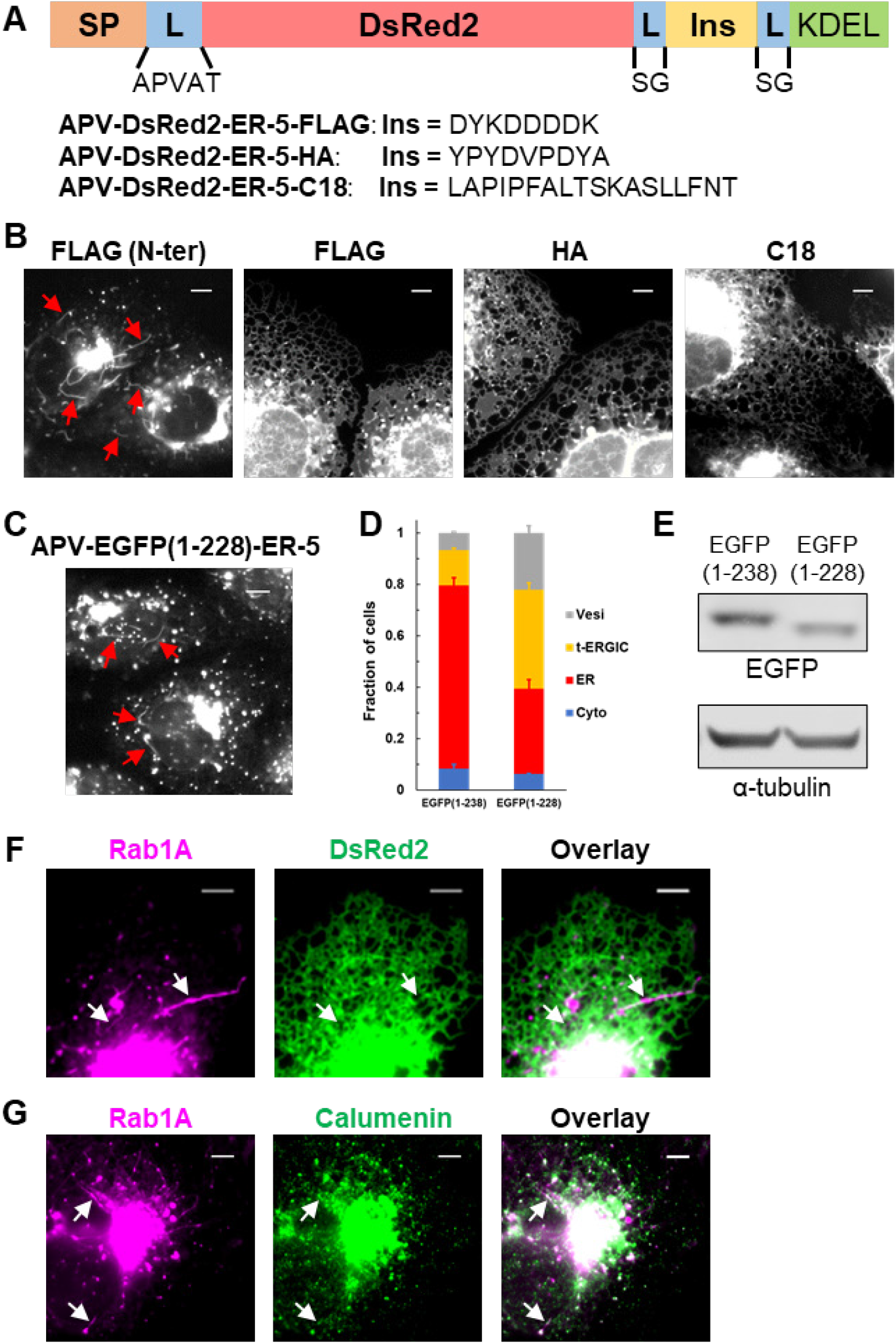
SURF4-KDELR Antagonism Determines the Localization of Cargo Proteins, Related to Figure 6. (A) Schematics of different C-terminal insertions of APV-DsRed2-ER-5. (B) Representative fluorescence micrographs of APV-FLAG-DsRed2-ER-5, APV-DsRed2-ER-5-FLAG, APV-DsRed2-ER-5-HA, and APV-DsRed2-ER-5-C18 in COS-7 cells. (C) Representative fluorescence micrograph of APV-EGFP(1-228)-ER-5 in COS-7 cells, showing t-ERGICs. (D,E) Subcellular distributions (D) and immunoblots (E) of APV-EGFP(1-238)-ER-5 and the C-terminally truncated APV-EGFP(1-228)-ER-5. Error bar: SEM (n = 3 with ~50 cells in each replicate). The APV-EGFP(1-238)-ER-5 subcellular distribution in (D) duplicates “APV-EGFP-ER-5” in Figure S6B. (F) Dual-color live-cell fluorescence micrographs of EGFP-Rab1A and EPV-DsRed2-ER-5-HA in a co-transfected COS-7 cell. (G) Immunofluorescence of endogenous calumenin vs. fluorescence micrograph of EGFP-Rab1A in a COS-7 cell. Scale bars: 5 μm. Arrows point to t-ERGICs.

## DESCRIPTION OF SUPPLEMENTAL VIDEOS

Video S1. DsRed2-ER-5 in fast-moving tubular organelles in a COS-7 cell, related to Figure 1C. Time 117.0 sec in the video corresponds to “0 sec” in Figure 1C. Arrows point to the Golgi.

Video S2. DsRed2-ER-5-containing t-ERGIC moving along microtubules in a COS-7 cell. Green: DsRed2-ER-5. Magenta: 2xmCherry-EMTB (microtubules).

Video S3. Left: RUSH of SBP-DsRed2-ER-5 in a COS-7 cell, related to Figure 2D. 80 μM biotin was added at time 0. The fluorescent cargo was redistributed from the ER to the Golgi via the t-ERGIC. Right: Another example of RUSH experiment, related to Figure S2J. The peripheral ER fluorescence was moved to the central Golgi as the t-ERGIC cycles between the two. Timestamp: min:sec.

Video S4. RUSH of TPV-CXCL9-mCherry-SBP (left) and EPV-CXCL9-mCherry-SBP (right) in COS-7 cells, related to Figure 3O. 80 μM biotin was added at time 0. The faster ER-to-Golgi trafficking of TPV-CXCL9-mCherry-SBP was accompanied by the presence of t-ERGICs.

Video S5. Co-labeling of the t-ERGIC by APV-DsRed2-ER-5 (green) and AcGFP1-SURF4 (magenta) in a COS-7 cell.

Video S6. RUSH of APV-SBP-DsRed2-ER-5 (yellow) with co-transfection of AcGFP1-SURF4 (cyan) and HaloTag-Sec23A (labeled by JF635, magenta) in a COS-7 cell. 80 μM biotin was added at time 0. The newly generated t-ERGICs were labeled by SURF4 but not Sec23A. Timestamp: min:sec.

Video S7. Co-clustering of APV-SBP-DsRed2-ER-5 (green) and AcGFP1-SURF4 (magenta) into gradually expanding domains upon cargo release in RUSH in a COS-7 cell, related to Figure 5D. 80 μM biotin was added at time 0. Timestamp: min:sec.

Video S8. Dissolution of the SURF4-cargo condensates by 1,6-hexanediol in a COS-7 cell, related to Figure 5G. 80 μM biotin was added 30 min before the addition of 3% 1,6-hexanediol at time 0. Individual condensates labeled by APV-SBP-DsRed2-ER-5 (green) and AcGFP1-SURF4 (magenta) dissolved at different times, but always concurrently for the two color channels. Shrinkage in cell size may be attributed to the osmotic pressure of hexanediol.

## REFERENCES

Allan, B.B., Moyer, B.D., and Balch, W.E. (2000). Rab1 recruitment of p115 into a cis-SNARE complex: Programming budding COPII vesicles for fusion. Science 289, 444–448.

Appenzeller-Herzog, C., and Hauri, H.P. (2006). The ER-Golgi intermediate compartment (ERGIC): In search of its identity and function. J. Cell Sci. 119, 2173–2183.

Bannykh, S.I., Rowe, T., and Balch, W.E. (1996). The organization of endoplasmic reticulum export complexes. J. Cell Biol. 135, 19–35.

Barlowe, C., and Helenius, A. (2016). Cargo capture and bulk flow in the early secretory pathway. Annu. Rev. Cell Dev. Biol. 32, 197–222.

Belden, W.J., and Barlowe, C. (2001). Role of Erv29p in collecting soluble secretory proteins into ER-derived transport vesicles. Science 294, 1528–1531.

Ben-Tekaya, H., Miura, K., Pepperkok, R., and Hauri, H.-P. (2005). Live imaging of bidirectional traffic from the ERGIC. J. Cell Sci. 118, 357–367.

Blum, R., Stephens, D.J., and Schulz, I. (2000). Lumenal targeted GFP, used as a marker of soluble cargo, visualises rapid ERGIC to Golgi traffic by a tubulo-vesicular network. J. Cell Sci. 113, 3151–3159.

Boncompain, G., Divoux, S., Gareil, N., De Forges, H., Lescure, A., Latreche, L., Mercanti, V., Jollivet, F., Raposo, G., and Perez, F. (2012). Synchronization of secretory protein traffic in populations of cells. Nat. Methods 9, 493–498.

Brandizzi, F., and Barlowe, C. (2013). Organization of the ER-Golgi interface for membrane traffic control. Nat. Rev. Mol. Cell Biol. 14, 382–392.

Bräuer, P., Parker, J.L., Gerondopoulos, A., Zimmermann, I., Seeger, M.A., Barr, F.A., and Newstead, S. (2019). Structural basis for pH-dependent retrieval of ER proteins from the Golgi by the KDEL receptor. Science 363, 1103–1107.

Casler, J.C., Papanikou, E., Barrero, J.J., and Glick, B.S. (2019). Maturation-driven transport and AP-1–dependent recycling of a secretory cargo in the Golgi. J. Cell Biol. 218, 1582–1601.

Casler, J.C., Zajac, A.L., Valbuena, F.M., Sparvoli, D., Jeyifous, O., Turkewitz, A.P., Horne-Badovinac, S., Green, W.N., and Glick, B.S. (2020). ESCargo: a regulatable fluorescent secretory cargo for diverse model organisms. Mol. Biol. Cell 31, 2892–2903.

Dancourt, J., and Barlowe, C. (2010). Protein sorting receptors in the early secretory pathway. Annu. Rev. Biochem. 79, 777–802.

Dempsey, G.T., Vaughan, J.C., Chen, K.H., Bates, M., and Zhuang, X. (2011). Evaluation of fluorophores for optimal performance in localization-based super-resolution imaging. Nat. Methods 8, 1027.

Emmer, B.T., Hesketh, G.G., Kotnik, E., Tang, V.T., Lascuna, P.J., Xiang, J., Gingras, A.C., Chen, X.W., and Ginsburg, D. (2018). The cargo receptor SURF4 promotes the efficient cellular secretion of PCSK9. Elife 7, e38839.

Feng, Z., Chen, X., Wu, X., and Zhang, M. (2019). Formation of biological condensates via phase separation: Characteristics, analytical methods, and physiological implications. J. Biol. Chem. 294, 14823–14835.

Gomez-Navarro, N., and Miller, E. (2016). Protein sorting at the ER-Golgi interface. J. Cell Biol. 215, 769–778.

Gorur, A., Yuan, L., Kenny, S.J., Baba, S., Xu, K., and Schekman, R. (2017). COPII-coated membranes function as transport carriers of intracellular procollagen I. J. Cell Biol. 216, 1745–1759.

Hauri, H.P., and Schweizer, A. (1992). The endoplasmic reticulum-Golgi intermediate compartment. Curr. Opin. Cell Biol. 4, 600–608.

Hauser, M., Yan, R., Li, W., Repina, N.A., Schaffer, D.V., and Xu, K. (2018). The spectrin-actin-based periodic cytoskeleton as a conserved nanoscale scaffold and ruler of the neural stem cell lineage. Cell Rep. 24, 1512–1522.

Honoré, B. (2009). The rapidly expanding CREC protein family: members, localization, function, and role in disease. Bioessays 31, 262–277.

Huang, B., Wang, W., Bates, M., and Zhuang, X. (2008). Three-dimensional super-resolution imaging by stochastic optical reconstruction microscopy. Science 319, 810–813.

Itzhak, D.N., Tyanova, S., Cox, J., and Borner, G.H.H. (2016). Global, quantitative and dynamic mapping of protein subcellular localization. Elife 5, e16950.

Klumperman, J., Schweizer, A., Clausen, H., Tang, B.L., Hong, W., Oorschot, V., and Hauri, H.P. (1998). The recycling pathway of protein ERGIC-53 and dynamics of the ER-Golgi intermediate compartment. J. Cell Sci. 111, 3411–3425.

Kroschwald, S., Maharana, S., and Simon, A. (2017). Hexanediol: a chemical probe to investigate the material properties of membrane-less compartments. Matters 3, e201702000010.

Kurokawa, K., and Nakano, A. (2019). The ER exit sites are specialized ER zones for the transport of cargo proteins from the ER to the Golgi apparatus. J. Biochem. 165, 109–114.

Lee, M.C.S., Miller, E.A., Goldberg, J., Orci, L., and Schekman, R. (2004). Bi-directional protein transport between the ER and Golgi. Annu. Rev. Cell Dev. Biol. 20, 87–123.

Malkus, P., Jiang, F., and Schekman, R. (2002). Concentrative sorting of secretory cargo proteins into COPII-coated vesicles. J. Cell Biol. 159, 915–921.

Marra, P., Maffucci, T., Daniele, T., Tullio, G. Di, Ikehara, Y., Chan, E.K.L., Luini, A., Beznoussenko, G., Mironov, A., and De Matteis, M.A. (2001). The GM130 and GRASP65 golgi proteins cycle through and define a subdomain of the intermediate compartment. Nat. Cell Biol. 3, 1101–1113.

Mironov, A.A., and Beznoussenko, G. V. (2019). Models of intracellular transport: Pros and cons. Front. Cell Dev. Biol. 7, 146.

Mironov, A., Beznoussenko, G. V., Trucco, A., Lupetti, P., Smith, J.D., Geerts, W.J.C., Koster, A.J., Burger, K.N.J., Martone, M.E., Deerinck, T.J., Ellisman, M.H., and Luini, A. (2003). ER-to-Golgi carriers arise through direct en bloc protrusion and multistage maturation of specialized ER exit domains. Dev. Cell 5, 583–594.

Mitrovic, S., Ben-Tekaya, H., Koegler, E., Gruenberg, J., and Hauri, H.-P. (2008). The cargo receptors Surf4, endoplasmic reticulum-Golgi intermediate compartment (ERGIC)-53, and p25 are required to maintain the architecture of ERGIC and Golgi. Mol. Biol. Cell 19, 1976–1990.

Moyer, B.D., Allan, B.B., and Balch, W.E. (2001). Rab1 interaction with a GM130 effector complex regulates COPII vesicle cis-Golgi tethering. Traffic 2, 268–276.

Munro, S., and Pelham, H.R.B. (1987). A C-terminal signal prevents secretion of luminal ER proteins. Cell 48, 899–907.

Otte, S., and Barlowe, C. (2004). Sorting signals can direct receptor-mediated export of soluble proteins into COPII vesicles. Nat. Cell Biol. 6, 1189–1194.

Plutner, H., Cox, A.D., Pind, S., Khosravi-Far, R., Bourne, J.R., Schwaninger, R., Der, C.J., and Balch, W.E. (1991). Rab1b regulates vesicular transport between the endoplasmic reticulum and successive Golgi compartments. J. Cell Biol. 115, 31–43.

Presley, J.F., Cole, N.B., Schroer, T.A., Hirschberg, K., Zaal, K.J.M., and Lippincott-Schwartz, J. (1997). ER-to-Golgi transport visualized in living cells. Nature 389, 81–85.

Raykhel, I., Alanen, H., Salo, K., Jurvansuu, J., Van, D.N., Latva-Ranta, M., and Ruddock, L. (2007). A molecular specificity code for the three mammalian KDEL receptors. J. Cell Biol. 179, 1193–1204.

Saegusa, K., Sato, M., Morooka, N., Hara, T., and Sato, K. (2018). SFT-4/Surf4 control ER export of soluble cargo proteins and participate in ER exit site organization. J. Cell Biol. 217, 2073–2085.

Sannerud, R., Marie, M., Nizak, C., Dale, H.A., Pernet-Gallay, K., Perez, F., Goud, B., and Saraste, J. (2006). Rab1 defines a novel pathway connecting the pre-Golgi intermediate compartment with the cell periphery. Mol. Biol. Cell 17, 1514–1526.

Saraste, J., and Marie, M. (2018). Intermediate compartment (IC): from pre-Golgi vacuoles to a semi-autonomous membrane system. Histochem. Cell Biol. 150, 407–430.

Saraste, J., and Svensson, K. (1991). Distribution of the intermediate elements operating in ER to Golgi transport. J. Cell Sci. 100, 415–430.

Schindelin, J., Arganda-Carreras, I., Frise, E., Kaynig, V., Longair, M., Pietzsch, T., Preibisch, S., Rueden, C., Saalfeld, S., and Schmid, B. (2012). Fiji: an open-source platform for biological-image analysis. Nat. Methods 9, 676–682.

Schweizer, A., Fransen, J.A.M., Bachi, T., Ginsel, L., and Hauri, H.P. (1988). Identification, by a monoclonal antibody, of a 53-kD protein associated with a tubulo-vesicular compartment at the cis-side of the Golgi apparatus. J. Cell Biol. 107, 1643–1653.

Shomron, O., Nevo-Yassaf, I., Aviad, T., Yaffe, Y., Zahavi, E.E., Dukhovny, A., Perlson, E., Brodsky, I., Yeheskel, A., Pasmanik-Chor, M., Mironov, A., Beznoussenko, G.V., Mironov, A.A., Sklan, E.H., Patterson, G.H., Yonemura, Y., Kaether, C., and Hirschberg, K. (2019). Uncoating of COPII from ER exit site membranes precedes cargo accumulation and membrane fission. BioRxiv 727107.

Simpson, J.C., Nilsson, T., and Pepperkok, R. (2005). Biogenesis of tubular ER-to-Golgi transport intermediates. Mol. Biol. Cell 17, 723–737.

Stenmark, H. (2009). Rab GTPases as coordinators of vesicle traffic. Nat. Rev. Mol. Cell Biol. 10, 513–525.

Stephens, D.J. (2012). Functional coupling of microtubules to membranes - implications for membrane structure and dynamics. J. Cell Sci. 125, 2795–2804.

Stephens, D.J., Lin-Marq, N., Pagano, A., Pepperkok, R., and Paccaud, J.P. (2000). COPI-coated ER-to-Golgi transport complexes segregate from COPII in close proximity to ER exit sites. J. Cell Sci. 113, 2177–2185.

Tinevez, J.-Y., Perry, N., Schindelin, J., Hoopes, G.M., Reynolds, G.D., Laplantine, E., Bednarek, S.Y., Shorte, S.L., and Eliceiri, K.W. (2017). TrackMate: An open and extensible platform for single-particle tracking. Methods 115, 80–90.

Tisdale, E.J., Bourne, J.R., Khosravi-Far, R., Der, C.J., and Balch, W.E. (1992). GTP-binding mutants of Rab1 and Rab2 are potent inhibitors of vesicular transport from the endoplasmic reticulum to the golgi complex. J. Cell Biol. 119, 749–761.

Tsukumo, Y., Tsukahara, S., Saito, S., Tsuruo, T., and Tomida, A. (2009). A novel endoplasmic reticulum export signal: Proline at the +2-position from the signal peptide cleavage site. J. Biol. Chem. 284, 27500–27510.

Vorum, H., Hager, H., Christensen, B.M., Nielsen, S., and Honoré, B. (1999). Human calumenin localizes to the secretory pathway and is secreted to the medium. Exp. Cell Res. 248, 473–481.

Wang, J., Zhang, J., Lee, Y.M., Ng, S., Shi, Y., Hua, Z.-C., Lin, Q., and Shen, H.-M. (2017). Nonradioactive quantification of autophagic protein degradation with L-azidohomoalanine labeling. Nat. Protoc. 12, 279–288.

Wang, X., Wang, H., Xu, B., Huang, D., Nie, C., Pu, L., Zajac, G.J.M., Yan, H., Zhao, J., Shi, F., Emmer, B.T., Lu, J., Wang, R., Dong, X., Dai, J., Zhou, W., Wang, C., Gao, G., Wang, Y., Willer, C., Lu, X., Zhu, Y., and Chen, X.-W. (2021). Receptor-mediated ER export of lipoproteins controls lipid homeostasis in mice and humans. Cell Metab. 33, 350–366.

Westrate, L.M., Hoyer, M.J., Nash, M.J., and Voeltz, G.K. (2020). Vesicular and uncoated Rab1-dependent cargo carriers facilitate ER to Golgi transport. J. Cell Sci. 133, jcs239814.

Wilson, D.W., Lewis, M.J., and Pelhams, H.R.B. (1993). pH-dependent Binding of KDEL to Its Receptor. J. Biol. Chem. 268, 7465–7466.

Yan, R., Chen, K., and Xu, K. (2020). Probing nanoscale diffusional heterogeneities in cellular membranes through multidimensional single-molecule and super-resolution microscopy. J. Am. Chem. Soc. 142, 18866–18873.

Yin, Y., Garcia, M.R., Novak, A.J., Saunders, A.M., Ank, R.S., Nam, A.S., and Fisher, L.W. (2018). Surf4 (Erv29p) binds amino-terminal tripeptide motifs of soluble cargo proteins with different affinities, enabling prioritization of their exit from the endoplasmic reticulum. PLoS Biol. 16, e2005140.

Zanetti, G., Pahuja, K.B., Studer, S., Shim, S., and Schekman, R. (2012). COPII and the regulation of protein sorting in mammals. Nat. Cell Biol. 14, 20–28.

Zhao, Y.G., and Zhang, H. (2020). Phase separation in membrane biology: the interplay between membrane-bound organelles and membraneless condensates. Dev. Cell 55, 30–44.

